# Evidence for dual targeting of Arabidopsis plastidial glucose-6-phosphate transporter GPT1 to peroxisomes via the ER

**DOI:** 10.1101/2019.12.11.873000

**Authors:** Marie-Christin Baune, Hannes Lansing, Kerstin Fischer, Tanja Meyer, Lennart Charton, Nicole Linka, Antje von Schaewen

**Author notes:** Corresponding author: Antje von Schaewen. The author responsible for distribution of materials integral to the findings presented in this article in accordance with the policy described in the Instructions for Authors (www.plantcell.org) are Antje von Schaewen and Nicole Linka.

## Abstract

Former studies on Arabidopsis glucose-6-phosphate/phosphate translocator isoforms GPT1 and GPT2 reported viability of *gpt2* mutants, however an essential function for GPT1, manifesting as a variety of *gpt1* defects in the heterozygous state during fertilization/seed set. Among other functions, GPT1 is important for pollen and embryo-sac development. Since previous work on enzymes of the oxidative pentose phosphate pathway (OPPP) revealed comparable effects, we investigated whether GPT1 might dually localize to plastids and peroxisomes. In reporter fusions, GPT2 was found at plastids, but GPT1 also at the endoplasmic reticulum (ER) and around peroxisomes. GPT1 contacted oxidoreductases and also peroxins that mediate import of peroxisomal membrane proteins from the ER, hinting at dual localization. Reconstitution in yeast proteoliposomes revealed that GPT1 preferentially exchanges glucose-6-phosphate for ribulose-5-phosphate. Complementation analyses of heterozygous *gpt1* plants demonstrated that GPT2 is unable to compensate for GPT1 in plastids, whereas genomic *GPT1* without transit peptide (enforcing ER/peroxisomal localization) increased *gpt1* transmission significantly. Since OPPP activity in peroxisomes is essential during fertilization, and immuno-blot analyses hinted at unprocessed GPT1-specific bands, our findings suggest that GPT1 is indispensable at both plastids and peroxisomes. Together with the G6P-Ru5P exchange preference, dual targeting explains why GPT1 exerts functions distinct from GPT2 in Arabidopsis.

**One sentence summary:** In contrast to plastidial GPT2, GPT1 exhibits slightly different exchange preferences and alternatively targets the ER, from where the protein can be relocated to peroxisomes on demand.

## INTRODUCTION

In plant cells, the oxidative pentose phosphate pathway (OPPP) is found in plastids and the cytosol (reviewed in Kruger and von Schaewen, 2003), but transiently also in peroxisomes (Meyer et al., 2011; Hölscher et al., 2014; 2016). In each subcellular compartment, the OPPP has distinctive functions and thus requires subcellular distribution of the corresponding enzymes and their metabolites.

During the day, NADPH is provided by photosynthetic electron flow to ferredoxin-(Fd) NADP^+^ oxidoreductase (FNR; Palatnik et al., 2003), whereas at night, the OPPP is the main source of NADPH in chloroplasts and in heterotrophic plastids of non-green tissues (Dennis et al., 1997). The oxidation of 1 mole glucose-6-phosphate (G6P) to ribulose-5-phosphate (Ru5P) produces 2 moles of NADPH (at the expense of CO_2_ release) in three enzymatic steps: i) glucose-6-phosphate dehydrogenase (G6PD), ii) 6-phosphogluconolactonase (6PGL), and iii) 6-phospho-gluconate dehydrogenase (6PGD). These irreversible reactions are followed by reversible OPPP steps in the stroma, comprising transketolase (TK) and transaldolase (TA) that create a broad range of phosphorylated intermediates. Since the reversible OPPP reactions share intermediates with the Calvin cycle, they are essential for plant metabolism (reviewed in Kruger and von Schaewen, 2003). In the cytosol of plant cells only the irreversible OPPP reactions occur (Schnarrenberger et al., 1995), linked to the full cycle in plastids via epimerization of Ru5P to Xu5P and import by the Xylulose-5-phosphate/phosphate translocator (XPT) in the inner envelope (Eicks et al., 2002).

NADPH is the preferred reducing equivalent of anabolic reactions, both in plastids and the cytosol, needed mostly for the biosynthesis of amino acids, fatty acids, and nucleotides (Hutchings et al., 2005; Geigenberger et al., 2005). Furthermore, NADPH is important for redox homeostasis of the glutathione pool (GSH/GSSG) via NADPH-dependent glutathione-disulfide reductases (GRs). Arabidopsis GR1 dually localizes in the cytosol and peroxisomes (Marty et al., 2009; Mhamdi et al., 2010; Kataya and Reumann, 2010) and GR2 in plastids and mitochondria (Marty et al., 2019). Hence, OPPP reactions play an important role in plant cells (Kruger and von Schaewen, 2003), particularly with the onset of stress or developmental change. Such conditions are often linked to physiological sink states induced by pathogen infection of leaves and related signaling. Resulting callose formation at plasmodesmata leads to sugar accumulation in the cytosol that stimulates G6PDH activity/expression and NADPH production via the OPPP (Hauschild and von Schaewen, 2003; Scharte et al., 2009; Stampfl et al., 2016). Concomitantly activated NADPH oxidases at the plasma membrane (in plants called respiratory burst oxidase homologues, Rbohs; Torres et al., 2002) use cytosolic NADPH for extrusion of reactive oxygen species (ROS) into the apoplast. Superoxide (O_2_) is converted to hydrogen peroxide (H_2_O_2_) that enters the cell via aquaporins, leading to redox signaling in the cytosol. H_2_O_2_ is dissipated by peroxiredoxins (Prx), which in turn retrieve electrons from glutaredoxins (Grx) and thioredoxins (Trx), and the resulting dithiol-disulfide changes modulate cognate target enzymes in a similar manner (reviewed in Noctor and Foyer, 2016; Waszczak et al., 2018; Liebthal et al., 2018). This scenario also accompanies abiotic stress responses (e.g. to drought or salt), together with phosphorylation cascades activated in parallel (Pitzschke et al., 2006; dal Santo et al., 2012; Fancy et al., 2016; Landi et al., 2016).

OPPP enzymes were also found in purified plant peroxisomes (Corpas et al., 1998; del Río et al., 2002; Reumann et al., 2007; Hölscher et al., 2016), where they may serve as NADPH source to establish redox homeostasis via dual cytosolic/peroxisomal GR1 (Kataya and Reumann, 2010). Besides, NADPH is needed for metabolic reactions that occur exclusively in peroxisomes, like removal of double bonds in unsaturated fatty acid/acyl chains prior to β-oxidation, which includes final steps of auxin/jasmonic acid biosynthesis (Reumann et al., 2004). We previously reported that dual targeting of *Arabidopsis thaliana* OPPP enzymes G6PD1 (At5g35790, OPPP step 1) and PGL3 (At5g24400, OPPP step 2) to plastids and peroxisomes depends on the cytosolic redox state (Meyer et al., 2011; Hölscher et al., 2014). Furthermore, plants heterozygous for peroxisomal isoform PGD2 (At3g02360, OPPP step 3) failed to produce homozygous offspring due to mutual sterility of the *pgd2* gametophytes. This indicated for the first time an essential function of the OPPP in peroxisomes (Hölscher et al., 2016).

OPPP activity in organelles requires flux of intermediates across the corresponding membranes. In Arabidopsis, G6P import into plastids involves G6P/phosphate translocator GPT1 (At5g54800) and GPT2 (At1g61800) in the inner envelope membrane (Kammerer et al., 1998; Eicks et al., 2002; Knappe et al., 2003; Niewiadomski et al., 2005). In case of peroxisomes, phosphorylated metabolites with a huge hydration shell are likely unable to pass the porin-like channel described for malate and oxaloacetate (134 and 130 Da) first described in spinach (Reumann et al., 1996). In mammalian cells, Rokka et al. (2009) measured that only molecules below 200 Da are able to pass the pore-like channel of Pxmp2. G6P and Ru5P/Xu5P are larger (258 Da and 230 Da), implying that they are unlikely transported via peroxisomal porins. Thus, the issue of OPPP substrate and product transport across peroxisomal membranes remained unclear so far.

To provide the peroxisomal OPPP reactions with substrate, we reasoned that one of the two Arabidopsis GPT proteins may dually localize to peroxisomes, similar to originally plastid-annotated OPPP isoforms G6PD1 (Meyer et al., 2011) and PGL3 (Kruger and von Schaewen, 2003; Reumann et al., 2004; Hölscher et al., 2014). GPT1 and GPT2 show 81% identity at the amino-acid level and catalyze the import of G6P into heterotrophic plastids needed for starch synthesis and NADPH provision via the stromal OPPP reactions (Kammerer et al., 1998). GPT2 expression is most abundant in heterotrophic tissues (senescing leaves, sepals, seeds) and can be induced by high light in leaves (Athanasiou et al., 2010; Weise et al., 2019), whereas GPT1 is ubiquitously expressed, with highest levels in reproductive tissues (Niewiadomski et al., 2005; Kunz et al., 2010). Loss of GPT2 function reduced starch levels, but yielded vital plants (Niewiadomski et al., 2005; Kunz et al., 2010; Athanasiou et al., 2010; Dyson et al., 2014; 2015). However, lack of GPT1 was detrimental, leading to an early arrest of pollen and ovule development. Resulting gametophyte and embryo lethality showed as incompletely filled siliques (Niewiadomski et al., 2005; Andriotis et al., 2010; Flügge et al., 2011).

We noticed that GPT1 displays a canonical C-terminal peroxisomal targeting signal type 1 (PTS1 motif AKL) that matches the consensus (S/A)-(K/R)-(L/M/I) of soluble proteins (Gould et al., 1989; Reumann, 2004; Platta and Erdmann, 2007; Reumann and Bartel, 2016). This seemed odd, since peroxisomal membrane proteins (PMPs) exhibit independent mPTS motifs of varying sequence (Rottensteiner et al., 2004). In general, two classes of PMPs are known. Class-I PMPs are directly inserted into peroxisomal membranes (PerMs) from the cytosol, which involves peroxins Pex3 and Pex19 (in some organisms also Pex16; Platta and Erdmann, 2007). By contrast, class-II PMPs are first inserted into the endoplasmic reticulum (ER) via the Sec61 import pore and then transported to the peroxisomal ER (perER), from where peroxisomes are formed *de novo* (Theodoulou et al., 2013; Reumann and Bartel, 2016; Kao et al., 2018). The exact mechanism remains to be resolved, but involvement of Pex16 and Pex3 for ER recruitment and sorting to peroxisomes is most likely (Aranovich et al., 2014). Interestingly, mutation of Arabidopsis *PEX16* resulted in a shrunken seed phenotype (*sse1*) with impaired fatty acid biosynthesis (Lin et al., 1999, 2004), reminiscent of some *gpt1* defects (Niewiadomski et al., 2005), but no defects in pollen germination.

Here we report that both GPT1 and GPT2 may insert into the ER, but only the N-terminal part of GPT1 is able to initiate ER targeting, a prerequisite shared with class-II PMPs. Co-expression of various reporter fusions was used to analyze subcellular localization and protein interaction of GPT1 in plant cells. GPT1 formed homodimers at plastids, but not readily at the ER, and interacted with two cytosolic oxidoreductases listed by the Membrane-based Interactome Network Database (MIND) for Arabidopsis proteins with 38% confidence (Lalonde et al., 2010; Chen et al., 2012; Jones et al., 2014). In addition, we found evidence for transient interaction of GPT1 with early peroxins involved in PMP delivery via the ER. As rare event, GPT1-reporter fusions were detected in membrane structures surrounding peroxisomes. Our main questions were: 1) which protein parts confer dual targeting; 2) how this may be regulated; 3) which OPPP metabolite leaves peroxisomes; and 4) whether some defects of heterozygous *gpt1* mutant plants (Niewiadomski et al., 2005) may be related to missing transport across peroxisomal membranes during fertilization.

## RESULTS

### GPT1 dually targets plastids and the ER

The alignment of GPT1 and GPT2 protein sequences from different *Brassicaceae* (Supplemental Figure 1) revealed that the isoforms mostly diverge at their N-terminal ends, whereas the central transmembrane regions (for substrate binding/transport) are highly conserved. Subcellular targeting was studied with various N- and C-terminal reporter fusions of the two Arabidopsis GPT isoforms and examined in transfected protoplasts (Arabidopsis or tobacco) by confocal laser-scanning microscopy (CLSM).

All N-terminally masked/truncated GPT variants (Supplemental Figure 2A) localized at the ER (Supplemental Figure 2B, green signals) as determined by co-expression with organelle markers (magenta signals), i.e. G/OFP-ER (Rips et al., 2014) or peroxisome (Per) marker G/OFP-PGL3_*C-short* (formerly named G/OFP-PGL3(∼50aa)-SKL; Meyer et al., 2011). Note that co-localization of green and magenta signals appears white. Both GPT fusions occasionally formed *Z membranes* (Supplemental Figure 2B, white patches), a term coined for overexpressed integral membrane proteins (Gong et al., 1996). GPT1_*C-full* labeled ring-like substructures of the ER, approximately 3 µm in diameter (Supplemental Figure 2C, panel b), and interfered with import of the peroxisome marker (Supplemental Figure 2B, panel n), which was never observed for GPT2_*C-full* (Supplemental Figure 2B, panel p). Mutagenesis of GPT1-AKL to -AKQ (or GPT2-AKQ to -AKL) had no effect on localization of the fusion proteins (not shown).

Among the C-terminal reporter fusions, localization of GPT1 also differed from GPT2. The GPT1 *full-length* version (Figure 1A), with GFP pointing to the plastid stroma (or cytosol, when inserted into the ER), was spotted at both plastids and the ER (Figure 1B, panels a,c, arrowheads), but GPT2 only at plastids (Figure 1B, panels b,d, green signals; for single channel images, see Supplemental Figure 3B). A region comprising the N-terminus plus first five membrane domains (*N-5MD*, 1-240 amino acids) with OFP pointing to the intermembrane space (IMS), labeled the plastid surface (Supplemental Figure 4B, panels a-d; green signals). The N-terminus plus first two membrane domains (*N-2MD,* 1-155 amino acids) with GFP pointing to the stroma showed patchy plastid labeling, indicative of partial reporter cleavage (Supplemental Figure 4B, panels e-h), and in case of GPT1 also ER labeling (Figure 4B, panels e, and f, arrowheads), albeit to varying extent (Supplemental Figure 4C, panels a-e). Again, small ring-like structures of peroxisomal size were labeled by GFP, but without surrounding the peroxisome marker (Supplemental Figure 4C, panel e, single sections). With the N-terminal region (*N-term*, 1-91/92 amino acids) fused to the reporter, stroma labeling was observed for both GPT proteins (Supplemental Figure 4B, panels i-l). These results indicated that the region comprising the N-terminus plus first two transmembrane GPT1 domains is important for alternative targeting to the ER.

**Figure 1.**
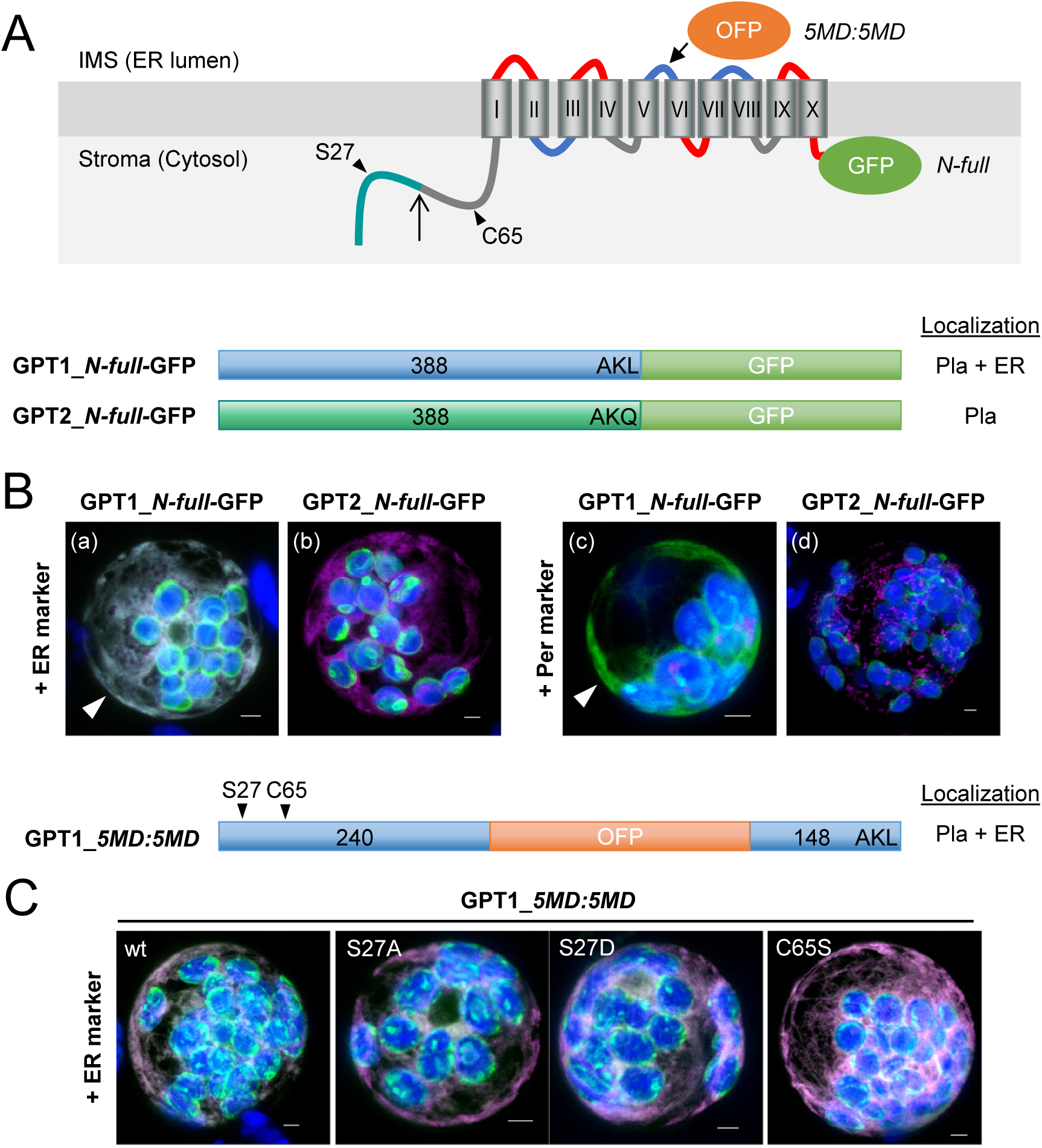
GPT1 reporter fusions dually localize to plastids and the ER. **A**, Topology model of Arabidopsis glucose-6-phosphate/phosphate translocator (GPT) isoforms with 10 membrane domains (MD) depicted as barrels (roman numbering), connected by hinge regions (red, positive; blue, negative; grey, neutral net charge), and both N-/C-terminal ends facing the stroma (Lee et al. 2017). Relevant positions are indicated: Plastidic transit peptide (TP, green), TP processing site (upward arrow), N-terminal amino acids potentially modified/regulatory in GPT1 (arrowheads), medial OFP insertion (*5MD:5MD*) and C-terminal GFP fusion (*N-full*). ER, endoplasmic reticulum; IMS, intermembrane space. **B-C**, Localization of the depicted GPT-reporter fusions upon transient expression in Arabidopsis protoplasts (24-48 h post transfection). **B**, With free N-terminus, GPT1 targets both plastids and the ER (panels a and c, arrowheads), but GPT2 only plastids (Pla; panels b and d). **C**, The medial GPT1_*5MD:5MD* construct (wt, wildtype) was used for analyzing potential effects of single amino acid changes in the N-terminus: S27A (abolishing phosphorylation), S27D (phospho-mimic) and C65S (precluding S modification). All images show maximal projections of approximately 30 optical sections (Merge, for single channel images, see Supplemental Figure 5). Candidate fusions in green, ER marker (panel B, OFP-ER; panel C, GFP-ER) or peroxisome marker (Per; OFP-PGL3_C-*short*) in magenta, and chlorophyll fluorescence in blue. Co-localization of green and magenta (or very close signals less than 200 nm) appear white in the Merge of all channels. Bars = 3 μm.

### The first 155 amino acids of GPT1 are crucial for ER targeting

To exclude localization artifacts by masking N- or C-terminal targeting signals, we also cloned GPT-fusions with internal reporter at two different positions (Supplemental Figure 5A). Again, the GPT1 versions (GPT1_*2MD:8MD* and GPT1_*5MD:5MD*) labeled both plastids and the ER (Supplemental Figure 5B, panels a,b and e,f; arrowheads), whereas the GPT2 versions (GPT2_*2MD:8MD* and GPT2_*5MD:5MD*) only plastids (Supplemental Figure 5B, panels c,d and g,h). Protoplasts expressing the GPT_*2MD:8MD* fusions were additionally treated with Brefeldin A (BFA), which interfered with delivery of peroxisomal ascorbate peroxidase (pxAPX) via the ER (Mullen et al., 1999). BFA treatment abolished GPT1 signals at the ER, but not at plastids (neither of GPT2; Supplemental Figure 6). This confirmed direct GPT targeting to plastids, and that only GPT1 may insert into the ER.

Since alternative GPT1 localization seemed mediated by the soluble N-terminal part that strongly differs from GPT2 (Figure S1), amino acid positions suspected to be subject to post-translational modification were changed by site-directed mutagenesis in the medial GPT1_*5MD:5MD* fusion (Figure 1C). However, neither S27 (listed by PhosPhAt 4.0; Zulawski et al., 2013) changed to alanine (A, abolishing phosphorylation) or aspartate (D, mimicking phosphorylation; Ackerley et al., 2003), nor single cysteine C65 changed to serine (S, precluding redox modification) interfered with ER targeting. Domain swaps among the corresponding unmodified *medial* reporter constructs (Figure 2A) resulted in dual localization of GPT1_*2MD:8MD*_GPT2 and GPT1_*5MD:5MD*_GPT2 to plastids and the ER (Figure 2B, panels a,b and e,f; arrowheads), but solely plastid localization of GPT2_*2MD:8MD*_GPT1 and GPT2_*5MD:5MD*_GPT1 (Figure 2B, panels e,d and g,h; for single channel images, see Supplemental Figure 7). These results proved that the GPT1 N-terminus (plus first two MDs) is crucial for initiating alternative ER targeting.

**Figure 2.**
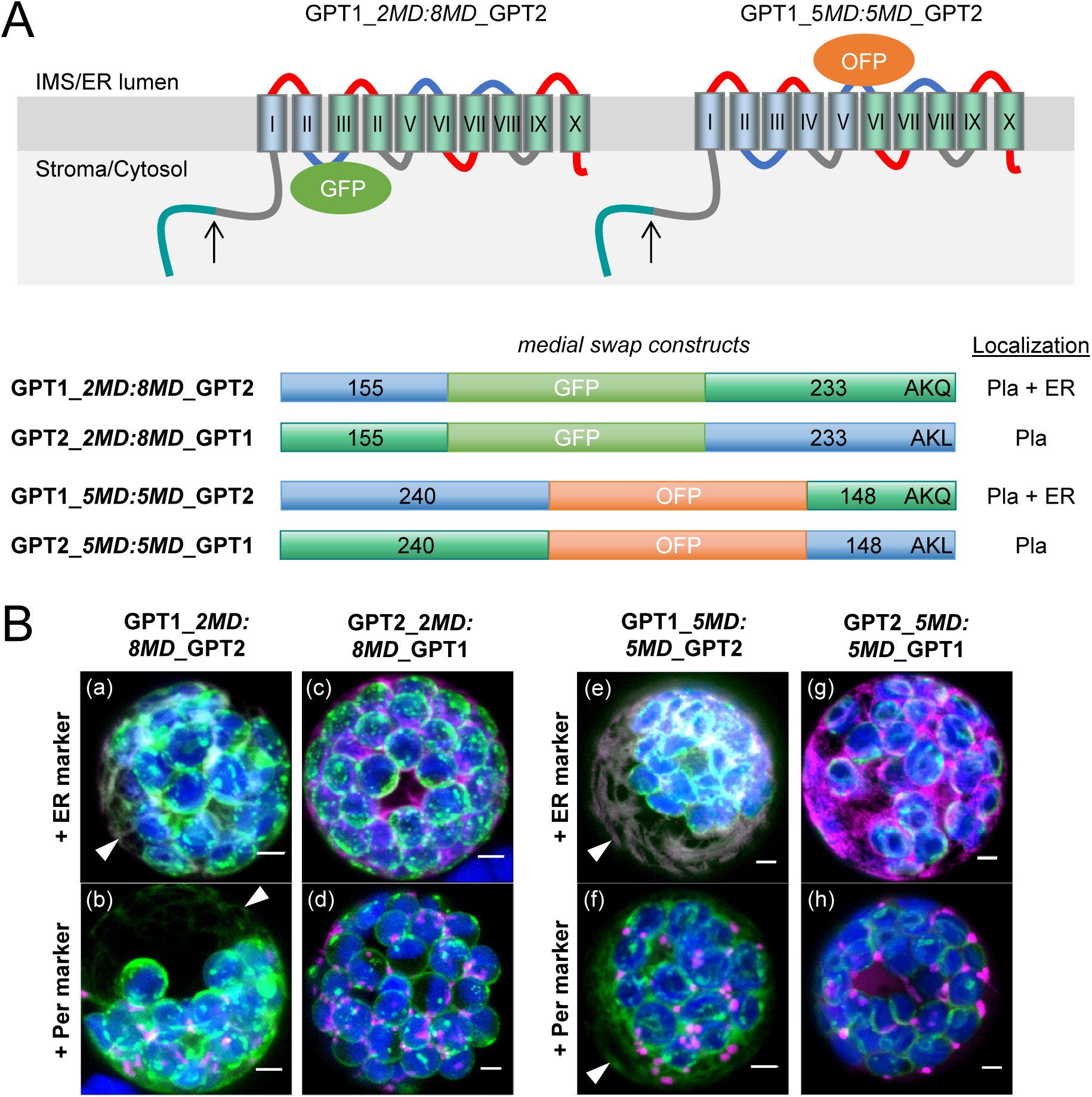
Domain swaps demonstrate that the N-terminus of GPT1 confers ER targeting. **A**, Topology models of the GPT *medial* swap constructs, with orientation of the inserted reporters: GFP facing the stroma/cytosol and OFP the intermembrane space (IMS)/lumen of the endoplasmic reticulum (ER). Membrane domains (depicted as barrels, roman numbering) of GPT1 in blue and of GPT2 in green. The upward arrows indicate transit peptide cleavage sites (plastid stroma). **B**, Localization of the indicated medial swap constructs in Arabidopsis protoplasts (24-48 h post transfection). When headed by GPT1 (GPT1_*2MD:8MD*_GPT2 or GPT1_*5MD:5MD*_GPT2), plastids and the ER (arrowheads) are labeled (panels a,b and e,f); when headed by GPT2 (GPT2_*2MD:8MD*_GPT1 or GPT2_*5MD:5MD*_GPT1), only plastids (Pla) are labeled (panels c,d and g,h). All images show maximal projections of approximately 30 optical sections (Merge, for single channel images, see Supplemental Figure 7). Candidate fusions in green, ER marker (G/OFP-ER) or peroxisome marker (Per; G/OFP-PGL3_C-*short*) in magenta, and chlorophyll fluorescence in blue. Co-localization of green and magenta (and very close signals less than 200 nm) appear white in the Merge of all channels. Bars = 3 μm.

### GPT1 dimer formation occurs at plastids and substructures of the ER

In functional form, the plastidial phosphate translocators are dimers composed of two identical subunits (Knappe et al., 2003). We therefore reasoned, if not necessary for ER targeting, amino acids S27 and/or C65 may be important for preventing GPT1 dimerization prior to reaching the final location(s). Therefore N- and C-terminal split YFP constructs of GPT1 were cloned and above described amino-acid changes introduced. Arabidopsis protoplasts were transfected and analyzed for GPT1-dimer formation (Figure 3) by bimolecular fluorescence complementation (BiFC; Walter et al., 2004). Reconstitution of the GPT1-split YFP combinations was detected only at plastids (Figure 3B, panels a-d), without effect of the indicated amino acid changes. In case of the split YFP-GPT1 fusions (enforcing ER insertion), large signal accumulations in the ER (including perinuclear structures) were observed for most variants. This signal did not represent the usually observed ER pattern and even affected distribution of the ER marker (see Figures 1 and 2). Among the amino acid changes analyzed, only C65S had an effect, resulting in hollow spherical structures surrounding single peroxisomes (Figure 3C, arrowhead) compared to the wild-type situation or S27 changes (Figure 3B, compare panels f-g to panel i, arrowhead; for single channel images, see Supplemental Figure 8). Thus, ER insertion seems not to require posttranslational modification, but sorting to PerMs may be negatively regulated by C65 modification.

**Figure 3.**
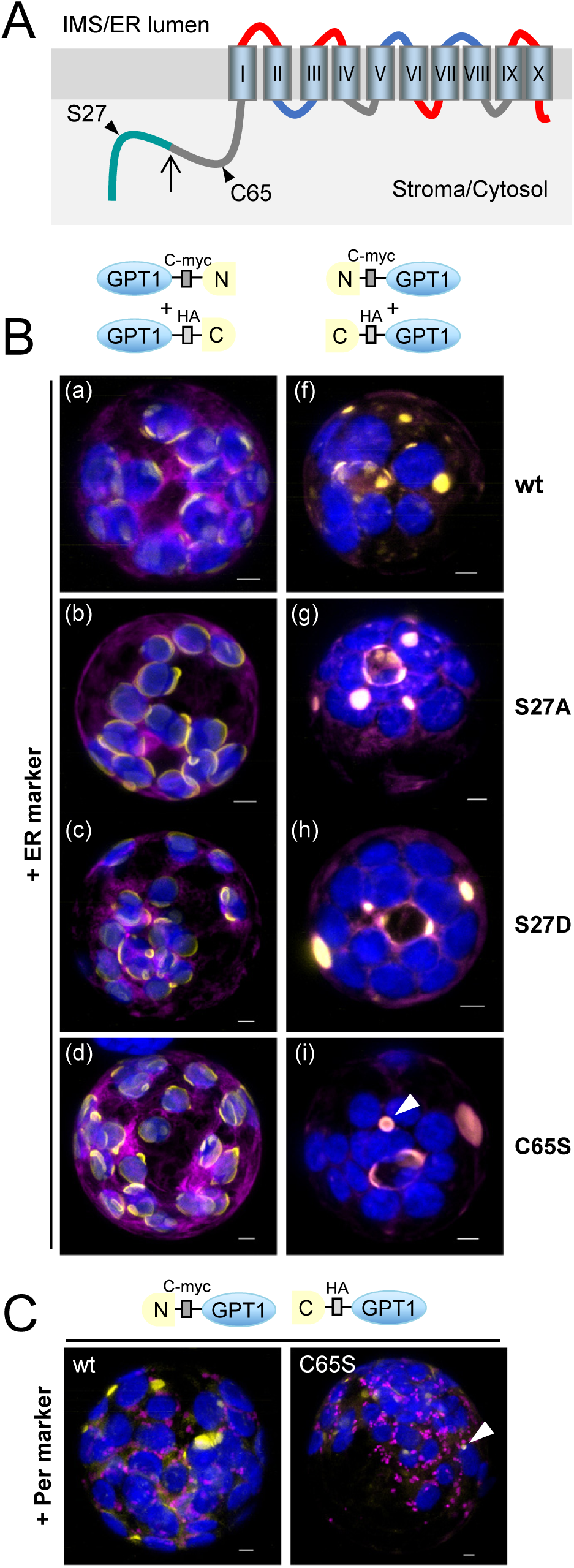
GPT1 dimer formation occurs at plastids and ER substructures. **A**, Topology model of GPT1 with N-terminal transit peptide (green) and cleavage site (upward arrow) plus position of amino acids S27 and C65 (arrowheads). The membrane domains are depicted as barrels (roman numbering) connected by hinge regions of different net charge (red, positive; blue, negative; grey, neutral). **B**, Localization of yellow BiFC signals (reconstituted split YFP, N+C halves) due to interaction of the GPT1 parts in Arabidopsis protoplasts (24-48 h post transfection). With unmasked N-terminus, GPT1 may label plastids and the ER (left panels), but with masked N-terminus only the ER (right panels). In addition to unmodified GPT1 wild-type (wt), mutant combinations S27A (non-phosphorylated), S27D (phospho-mimic) and C65S (precluding S modification) were analyzed. GPT1 dimer formation occurred at plastid rims (left panels) or ER substructures (right panels), with little impact of the S27 changes, but visible effect of C65S (hollow sphere in panel i; surrounding a peroxisome in C, arrowheads). Note that structures with BiFC signals on the right (panels f-i) are also labeled by the ER marker (most obvious in panel g). **C**, Localization of the indicated split YFP combinations in co-expression with the peroxisome (Per) marker. Note that in case of C65S, the ring-like BiFC signal surrounds a peroxisome (arrowhead). All images show maximal projections of approximately 30 optical sections (Merge; for single channel images, see Supplemental Figure 8). Organelle markers (OFP-ER or OFP-PGL3_*C-short*) in magenta, chlorophyll fluorescence in blue. Co-localization of yellow and magenta (or very close signals less than 200 nm) appear whitish in the Merge of all channels. Bars = 3 μm.

### GPT1 recruitment to the ER involves redox transmitters

To find potential interaction partners of GPT1, the Membrane-based Interactome Database (MIND) of Arabidopsis proteins (based on split ubiquitin reconstitution in yeast; Lalonde et al., 2010), was searched. Two cytosolic oxidoreductases, Thioredoxin *h7* (Trx*_h7_*, At1g59730) and Glutaredoxin *c1* (Grx*_c1_*, At5g63030), were among the 21 candidates listed with highest score (Supplemental Table 1). BiFC analyses in Arabidopsis protoplasts confirmed interaction of GPT1 with Trx*_h7_* (Figure 4A) and Grx*_c1_* (Figure 4B) at the ER and its substructures, but not at plastids (Figure 4A, panel b), and more clearly when the N-terminus of GPT1 was masked (enforcing ER insertion). Occasionally, ER-derived membranes around peroxisomes were labeled (Figure 4A, panel b and d; Figure 4B, panel b, arrowheads), which was less obvious when the N-terminus of Grx*_c1_* was masked by split YFP (Figure 4B, panels c,d). To enhance interaction among the Arabidopsis proteins, selected BiFC combinations were co-expressed with the other oxidoreductase as OFP fusion in heterologous tobacco protoplasts. Similar results were obtained (Figure 4C and D) and also smaller spherical structures (<3 µm) detected. Of note, in simple co-expression studies, both Trx*_h7_*-OFP and Grx*_c1_*-OFP partially overlapped with the ER marker (Supplemental Figure 9B, white signals), confirming predicted N-myristoylation, and co-localized with GPT1_*N-2MD*-GFP at the ER (Supplemental Figure 9C). These results are consistent with the two oxidoreductases assisting GPT1 insertion into the ER and/or sorting to peroxisomes.

**Figure 4.**
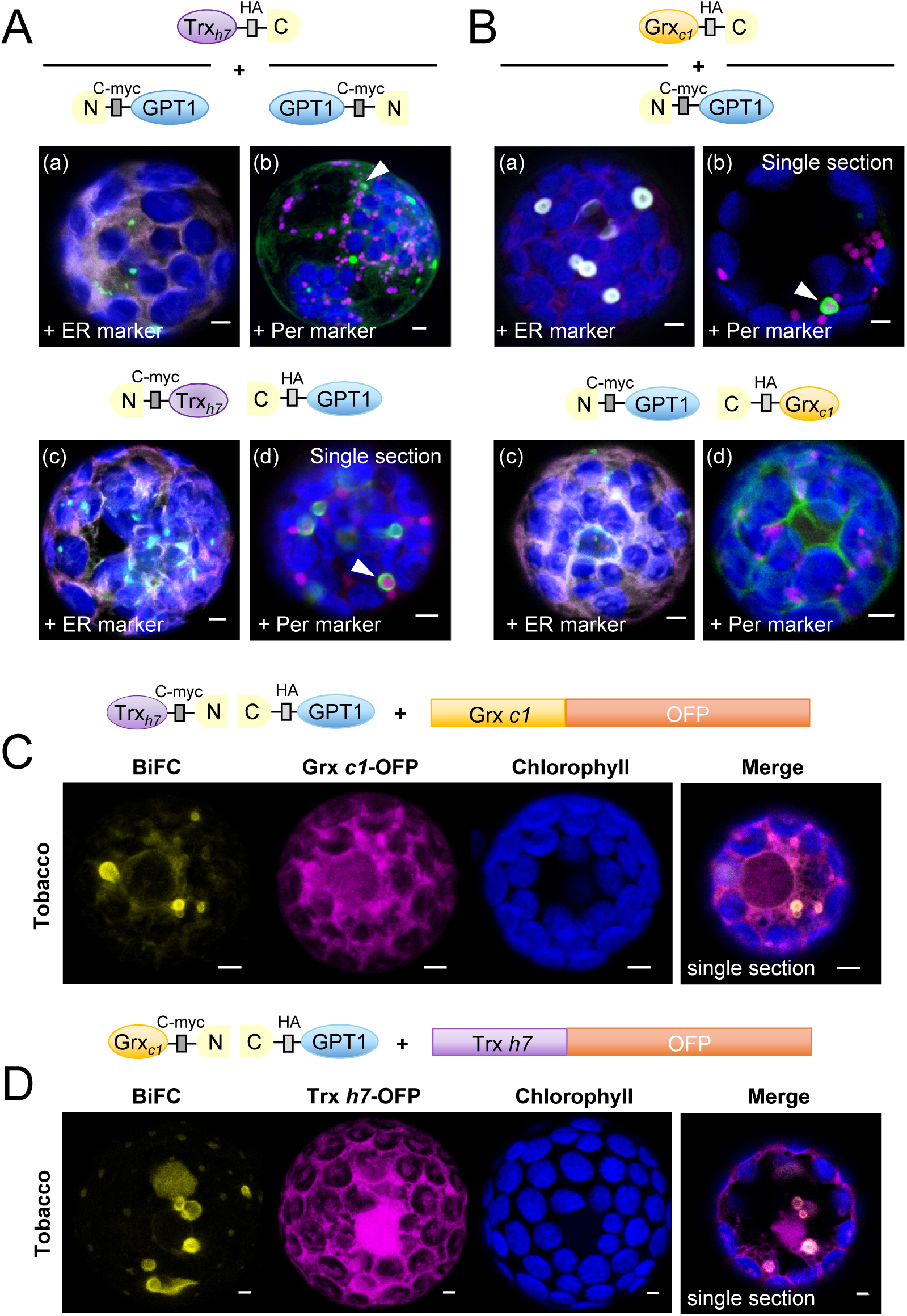
GPT1 interacts with cytosolic oxidoreductases Trx_h7_ and Grx_c1_ at the ER. **A-B**, Localization of GPT1 upon interaction with Trx h7 or Grx c1 in Arabidopsis protoplasts (24-48 h post transfection). The schemes illustrate different orientation of the candidate proteins with respect to free N- and C-terminal ends. GPT1 interacts with both oxidoreductases (green signals) at the endoplasmic reticulum (ER) and its spherical sub-structures (arrowheads), except when the N-terminus of Grx c1 is masked (B, panels c and d). Note that these substructures differ from those labelled in Figure 3B. Merge of BiFC signals (green) with ER marker (OFP-ER) or peroxisome marker (Per, OFP-PGL3_*C-short*) in magenta, and chlorophyll fluorescence in blue. **C-D**, Localization of split YFP reconstitution (BiFC, yellow signals) in heterologous tobacco protoplasts (24-48 h post transfection), testing a potential effect of the other oxidoreductase (co-expressed as OFP fusion, magenta). Note that similar ER substructures are labelled (Merge, single sections). All other images show maximal projections of approximately 30 optical sections. Chlorophyll fluorescence in blue. Co-localization and very close signals (less than 200 nm) appear white in the Merge of all channels. Bars = 3 μm.

### GPT1 contacts peroxins Pex3 and Pex16 at the ER

While class-I PMPs are inserted into PerMs directly from the cytosol (involving Pex3 and Pex19), class-II PMPs are first inserted into the ER (Platta and Erdmann, 2007). Since Pex3, Pex16, and Pex19 play also central roles during ER insertion, sorting of peroxisomal membrane proteins, and peroxisome biogenesis (Reumann and Bartel, 2016; Kao et al., 2018), we set out to analyze potential interaction with GPT1. In Arabidopsis, two Pex3 genes, Pex3-1 (At3g18160) and Pex3-2 (At1g48635; Hunt and Trelease, 2004), one Pex16 gene (At2g45690; Karnik and Trelease, 2005) and two Pex19 genes, Pex19-1 (At3g03490) and Pex19-2 (At5g17550; Hadden et al., 2006) exist. Analysis of N- and C-terminal reporter fusions in protoplasts revealed mainly PerM labeling for the two Pex3 isoforms, ER and PerM labeling for Pex16 (see also Lansing et al., 2019), and mostly cytosolic distribution for the two Pex19 isoforms (Supplemental Figure 10, shown for one of the two Pex3 and Pex19 isoforms). OFP-Pex3-1 displayed weak signals in the cytosol (not shown). BiFC analyses were conducted with Pex3-1, Pex16 and Pex19-1. GPT1 interaction with Pex3-1 and Pex16 was detected at PerMs, partially contiguous with the ER (Figure 5A, panels a,b). By contrast, GPT1 interaction with Pex19 was mostly distributed across the cytosol, but also labeled spherical structures (Figure 5A, panel d), when the C-terminal farnesylation motif (McDonnell et al., 2016) was accessible. Again, Pex16-GPT1 interaction interfered with import of the peroxisome (Per) marker (Figure 5A, panel b, magenta signals largely cytosolic), as already observed for GFP-GPT1_*C-full* (Supplemental Figure 2, panel n). Co-expression of GFP-GPT1_*C-full* with the OFP-based Pex fusions resulted in different patterns (Figure 5B), suggesting that the Pex interactions are merely transient. Co-expression with Pex3-1-OFP led in part to perinuclear localization of GFP-GPT1_*C-full*, reminiscent of the BiFC data obtained for GPT1 homodimerization (Figure 5B, panel a compared to Figure 3, panels f-i). Interestingly, Pex16 co-expression had visible effects on GPT1 localization, promoting concentration/vesiculation at the ER (Figure 5B, panel b), similar to Pex16 alone, but distinct from it (Supplemental Figure 10, compare B to C). In co-expression, Pex19-1 seemed to have no impact on GPT1 localization (Figure 5B, panels c and d).

**Figure 5.**
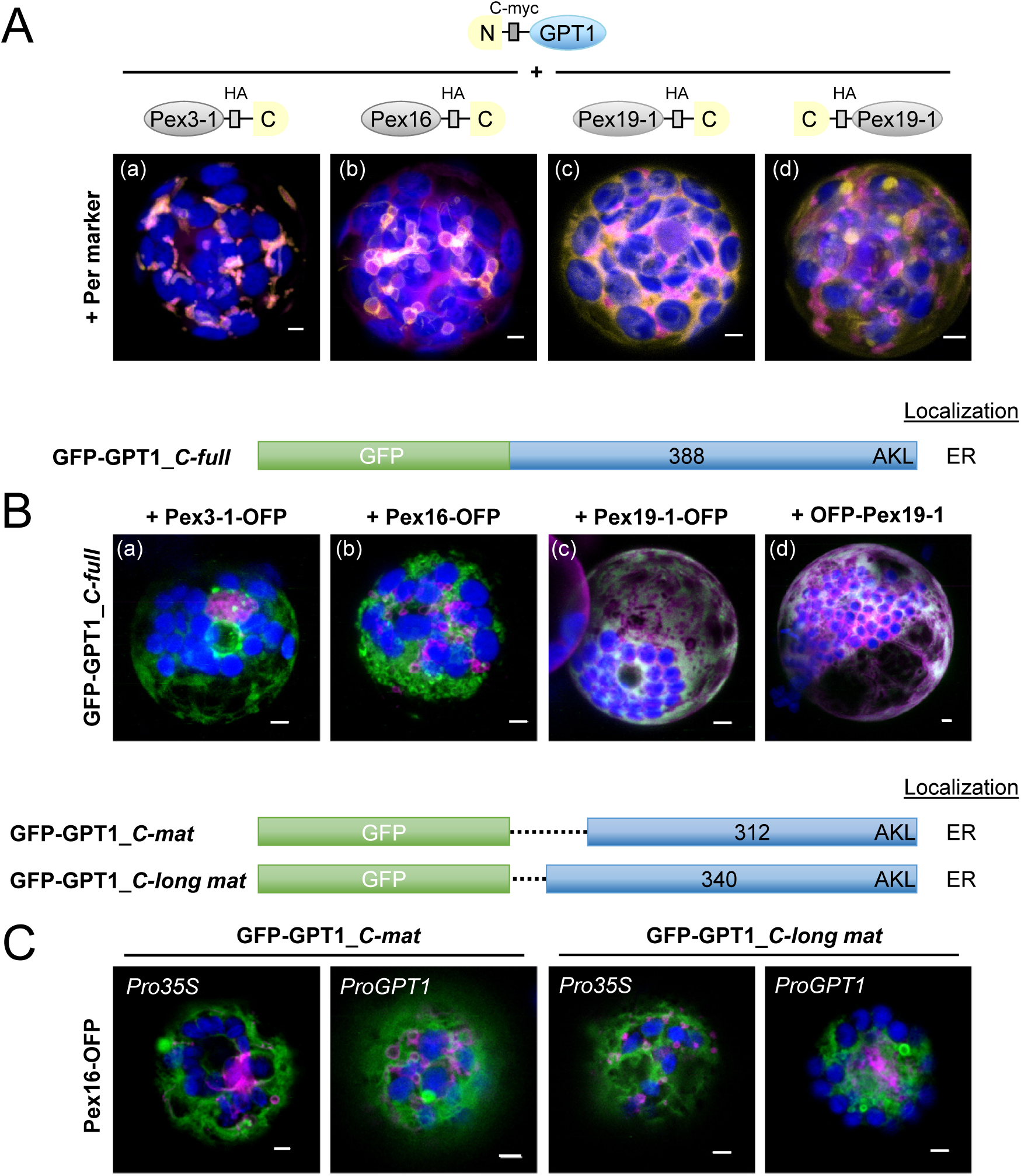
Interaction versus co-localization of GPT1 with Pex factors at the ER. **A**, Localization of the indicated split YFP combinations (yellow BiFC signals) in Arabidopsis protoplasts (24-48 h post transfection). Pex3, Pex16, and Pex19 are important for sorting a class of peroxisomal membrane proteins via the ER to peroxisomes. Per; soluble peroxisome marker (OFP-PGL3_*C-short*) in magenta. **B**, Co-expression of GFP-GPT1 and the corresponding Pex-OFP fusions indicates that interaction with the Pex factors is transient (isoforms Pex3-2 = At1g48635 and Pex19-2 = At5g17550 gave comparable results, not shown). Note that Pex16 co-expression has a vesiculating effect on GPT1 at the ER (Merge; for single channel images, see Supplemental Figure 10C). **A-B**, Maximal projections of approximately 30 optical sections. **C**, Co-expression of the indicated GFP-GPT1 fusions with Pex16-OFP in Arabidopsis protoplasts (72 h post transfection). The *C*_*mat* version lacks the entire N-terminal part (including C65), whereas *C_long mat* version lacks only the transit peptide (Supplemental Figure 1). Besides the 35S promoter (*Pro35S*), these GFP fusions were also expressed from the GPT1 promoter (*ProGPT1*), with similar results. Images show single optical sections (Merge; for single channel images, see Supplemental Figure 11). GFP fusions in green, Pex16-OFP in magenta and chlorophyll fluorescence in blue. Co-localization of green and magenta (or very close signals less than 200 nm) appear white in the Merge of all channels. Bars = 3 µm.

To make sure that the co-expression patterns obtained with Pex16 are no artifacts due to expression from the strong constitutive CaMV 35S promoter (*Pro35S*), two N-terminally truncated GPT1 versions (designated for stable plant transformation) were expressed also from the own promoter (*ProGPT1*), which gave comparable results (Figure 5C, for single channel images, see Supplemental Figure 11). Together with above BiFC analyses (Figure 5A), this demonstrated that ER-inserted GPT1 can be dragged to PerMs, and thus behaves like a class-II PMP that requires a special trigger to contact partner(s) (including Pex3 and Pex16) to reach mature peroxisomes.

### GPT1 may be recruited to peroxisomes and preferentially exchanges G6P for Ru5P

After plastid import, the N-terminal transit peptide (TP) of the precursor proteins is usually cleaved off (Schmidt et al., 1979; Chua and Schmidt, 1979). According to the recent elucidation of the 3-dimensional structure of the Arabidopsis triose-phosphate/phosphate translocator (Lee et al., 2017), both N- and C-terminal ends of GPT face the stroma. In case of GPT1 insertion into the ER, both the unprocessed N-terminus and C-terminal end should point to the cytosol, which was confirmed by topology analyses using *ro*GFP (Supplemental Figure 12). To test whether N-terminal modification or lack of transit-peptide processing might affect transport activity, we fused an N-terminal His tag (or GFP) to the *full-length* and *mature* GPT1 versions (with *mature* GPT2 as control) and measured metabolite exchange of the recombinant proteins in reconstituted yeast proteoliposomes (Linka et al., 2008). For the physiological exchange of G6P versus Pi using the mature versions (Figure 6A), His-matGPT1 reached about one third of the His-matGPT2 rates (with comparable expression levels in yeast cells, not shown). N-terminal modification by GFP did not affect the transport rates of GPT1, but presence of the transit peptide (equivalent to localization at the ER/PerMs) reduced transport rates by about half (not shown).

**Figure 6.**
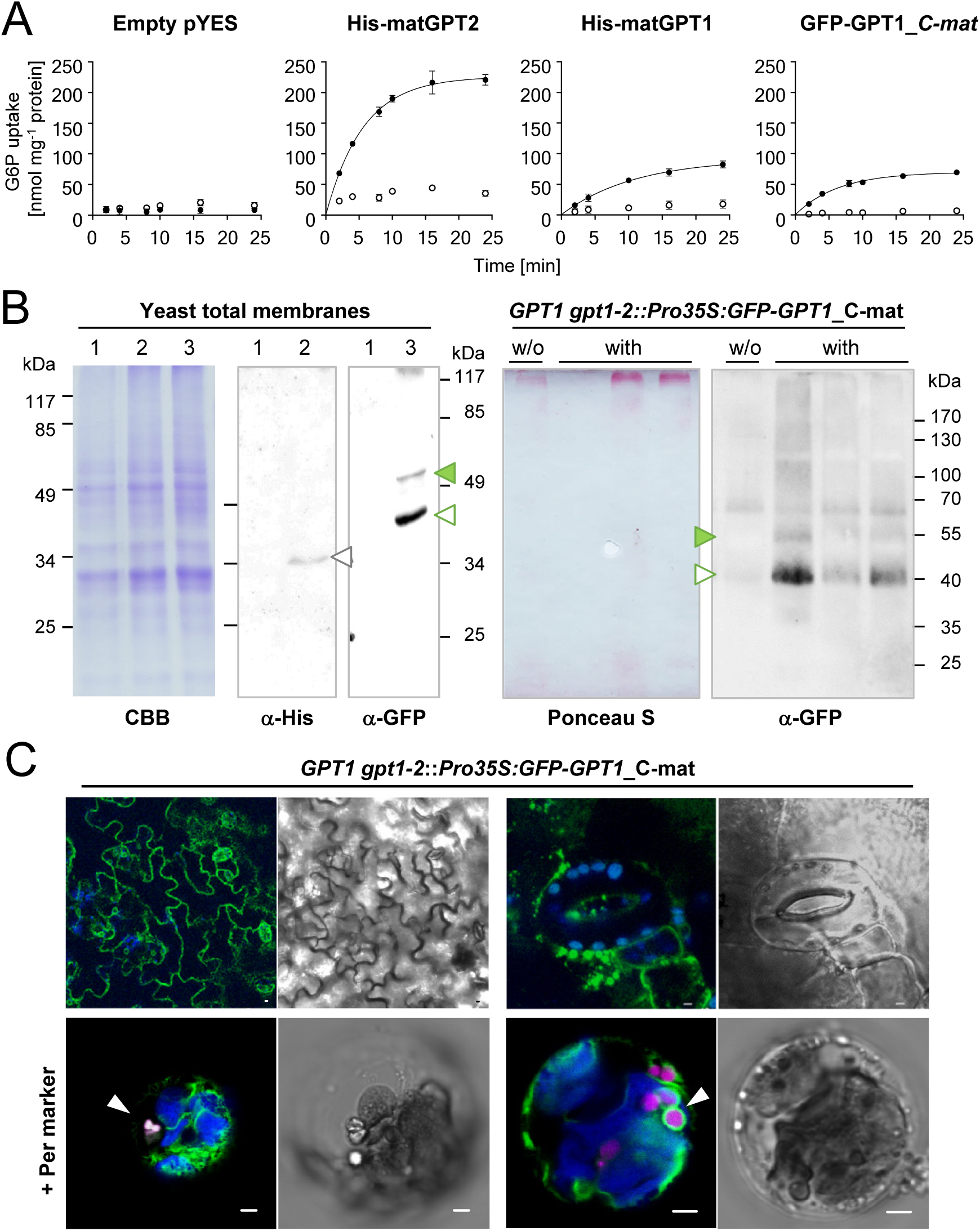
Transport activity and localization of mature GPT1 in yeast and plant cells. **A**, Time-dependent uptake of radioactively labeled [^14^C]-G6P (0.2 mM) into reconstituted proteoliposomes preloaded with 10 mM Pi (closed symbols) or without exchange substrate (open symbols) prepared from yeast cells harboring the empty vector (pYES) or the indicated GPT constructs. Note that transport rates of GPT1 are not influenced by the N-terminal tag (compare His-matGPT1 to GFP-matGPT1). In all graphs, the arithmetic mean of 3 technical replicates (±SD) was plotted against time (see Table 1 for substrate specificities). **B**, Immunoblot analysis upon expression in yeast and plant cells. Left, SDS gel of total yeast membrane fractions, stained with Coomassie brilliant blue (CBB) or blot detection by anti-His (α-His) or anti-GFP (α-GFP) antibodies: 1, empty vector; 2, His-matGPT1 (grey open triangle); 3, GFP-matGPT1 (green closed and open triangles). Right, blotted pellet fractions of leaf extracts (without detergent) prepared from Arabidopsis *GPT1 gpt1-2::Pro35S:GFP-GPT1_*C-mat plants (T2 progeny without (w/o) or with the transgene) developed with anti-GFP (α-GFP) antibodies. The Ponceau S-stained blot serves as loading reference. Note that GFP-GPT1 (closed green and open triangles) extracted from yeast or plant membranes migrate similarly. Bands of molecular masses are indicated (kDa). **C**, Localization of GFP-GPT1_*C-mat* in heterozygous *GPT1 gpt1-2* plants. Top, Green net-like structures (ER) in leaf epidermal cells (left), and spherical structures in seedlings (right); bars = 10 µm. Bottom, Pattern upon protoplast preparation and transfection with the peroxisome marker (Per; OFP-PGL3_*C-short*, magenta) in membranes surrounding peroxisomes (arrowheads). Chlorophyll fluorescence in blue. All images show single optical sections. Co-localization (and very close signals less than 200 nm) appear white in the Merge of all channels (bright field images shown as reference). Bars = 3 µm.

The *Pro35S:GFP-GPT1*_C-mat construct was stably introduced into heterozygous *gpt1-2* plants by floral dip transformation (Clough and Bent, 1998). Similar immunoblot patterns were obtained for the GFP-GPT1 proteins extracted from yeast or plant cells (Figure 6B, green arrowheads). In leaf cells of soil-grown plants, ER labeling dominated, but also spherical structures (≤3 µm) were detected (Figure 6C, top panels). Obviously, ER insertion of *mature* GPT1 occurs by default, but sorting to PerMs requires a stimulus. When mesophyll protoplasts were prepared from transgenic leaf material and transfected with the peroxisome (Per) marker (OFP-PGL3_*C-short*), GFP-labeled structures resembling newly forming peroxisomes appeared (Figure 6C, bottom panels; arrowheads).

If GPT1 imports G6P into peroxisomes, we wondered what might happen to Ru5P, the product of the three irreversible OPPP reactions. Especially, since analyses of the ribulose-5-phosphate epimerase (RPE) isoforms At1g63290 (cytosolic), At3g01850 (cytosolic), and At5g61410 (plastidic) (Kruger and Von Schaewen, 2003) did not give any hint on peroxisomal localization (unpublished data). We therefore analyzed, whether the *mature* GPT versions (with N-terminal His tag) may exchange G6P for Ru5P. As shown in Table 1, the relative velocity of matGPT1 was higher for G6P-Ru5P (112%) than for Pi-Ru5P exchange (59%), and differed from matGPT2 (87% for G6P-Ru5P and 75% for Pi-Ru5P). Importantly, exchange rates for 6-phosphogluconate (6PG <10%) were negligible.

**Table 1.** Initial velocities of Pi or G6P import for various exchange substrates. Time-dependent uptake of [^32^P]-Pi or [^14^C]-G6P (0.2 mM) into liposomes reconstituted with total yeast membranes of cells expressing the indicated *mature* GPT versions (nmol mg^-1^ total protein). Proteoliposomes were preloaded with 10 mM G6P, Ru5P, 6PG, or Pi. Relative velocities given in brackets were compared to the counter-exchange experiment Pi/G6P or G6P/Pi, which was set to 100%.

|  |  | His-matGPT1 | His-matGPT2 |
| --- | --- | --- | --- |
| **Pi versus** | **G6P** | **9.9** (100 %) | **19.3** (100 %) |
|  | **Ru5P** | **5.8** ( 59 %) | **14.4** ( 75 %) |
|  | **6PG** | **0.8** ( 8 %) | **1.2** ( 6 %) |
| **G6P versus** | **Pi** | **10.5** (100 %) | **32.6** (100 %) |
|  | **Ru5P** | **12.2** (116 %) | **28.3** ( 87 %) |
|  | **6PG** | **0.9** ( 9 %) | **3.1** ( 10 %) |

### Stress and developmental stimuli enhance ER targeting of GPT1

Since protoplast preparation (which is achieved by treating leaves with fungal enzymes) of stably transformed leaves led to recruitment of GFP-GPT1_*C-mat* to peroxisomes, we tested whether also treatment with a bacterial elicitor (flagellin) may affect GPT localization. Both, *GPT1-* and *GPT2-*N-full*-GFP* constructs were co-transfected with peroxisome (Per) marker OFP-PGL3_*C-short* in Arabidopsis protoplasts, samples were split in half, and analyzed after 24 h of mock or flg22 treatment. The latter led to enhanced GPT1 recruitment to the ER (Figure 7A, arrowheads), without major effect on plastid localization (GPT2 was neither affected; for single channel images, see Supplemental Figure 13).

**Figure 7.**
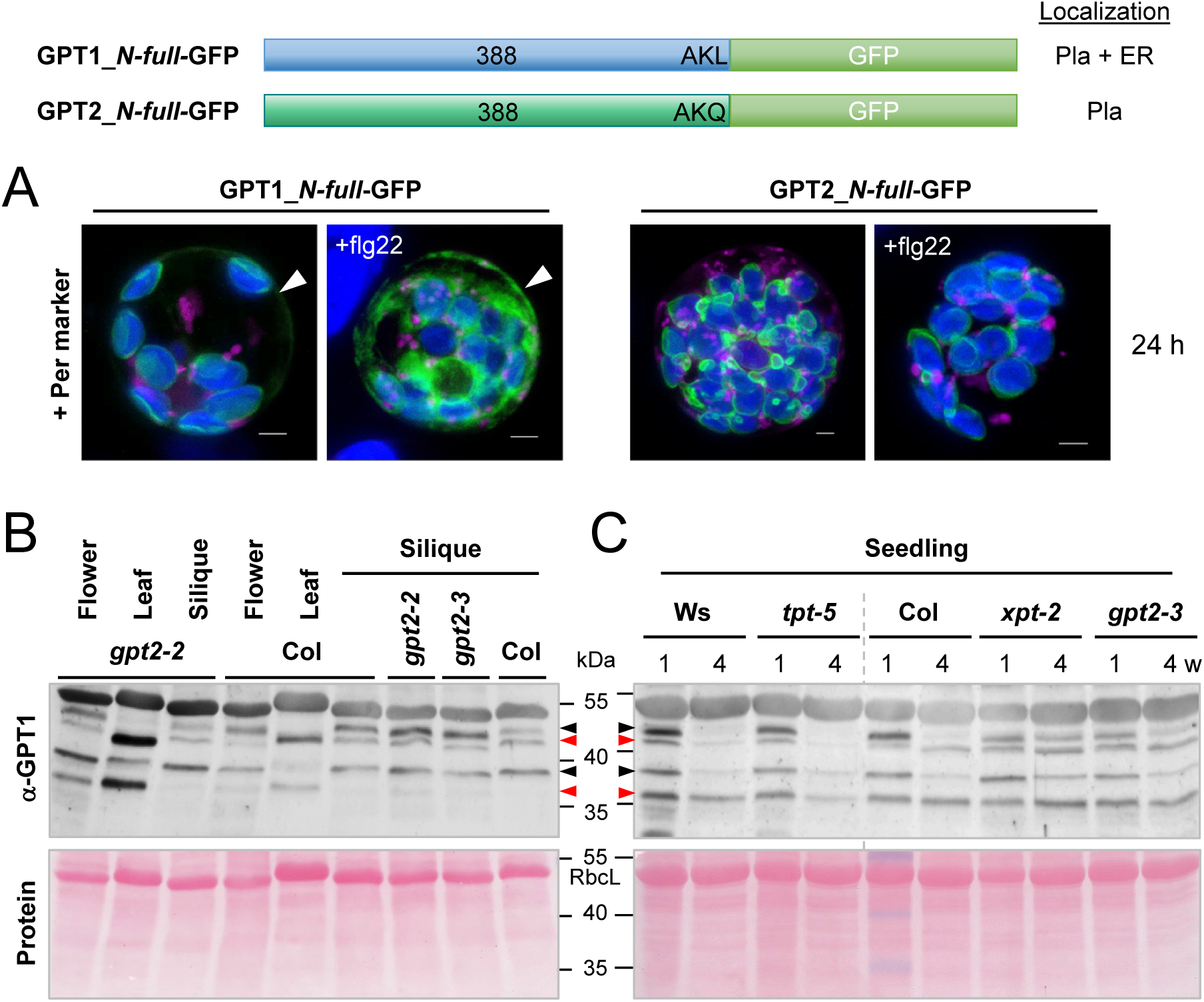
GPT1 detection at the ER is increased by stress treatment and in reproductive Arabidopsis tissues. **A**, Arabidopsis protoplasts were co-transfected with the indicated GPT-GFP fusions and the peroxisome marker (Per, OFP-PGL3_*C-short*), samples were split in half, one was treated with 0.2 µM flagellin peptide (+flg22), and the other mock-incubated for 24 h. Note that flg22 treatment did not change GPT localization to plastids, but enhanced the ER fraction of GPT1-GFP (arrowheads). All images show maximal projections of approximately 30 single sections (Merge; for single channel images, see Supplemental Figure S13). GFP fusions in green, peroxisome marker in magenta, and chlorophyll fluorescence in blue. Co-localization of magenta and green or very close signals (less than 200 nm) appear white in the Merge of all channels. Bars = 3 µm. **B-C**, Protein extracts (without detergent) of flower, leaf, and (green) silique tissue were prepared from wild-type plants (Col, Ws) and the indicated homozygous mutant lines. Supernatant fractions were separated on 10% SDS gels and blotted to nitrocellulose. After Ponceau-S staining, the blots were developed with GPT1-specific antibodies (α-GPT1) raised against the N-terminus with His-tag (Supplemental Figure S14). Arrowheads mark double bands of full-length GPT1 (predicted size: 42.3 kDa) and mature GPT1 (ca. 37-39 kDa, depending on TP processing). Red arrowheads point to bands suspected to represent a largely ‘off’ situation and black arrowheads the corresponding ‘on’ situation at either location (as deduced from comparison of leaf to silique tissue), likely due to protein modification. **C**, Immunoblot of seedlings harvested from germination plates (1% sucrose) after 1- or 4-week (w) growth in short-day regime. Included mutant alleles: *gpt2-2* (GK-950D09, T-DNA intron 2/exon 3), *gpt2-3* (GK-780F12, T-DNA in exon 4), *tpt-5* (FLAG_124C02, T-DNA in exon 9), and *xpt-2* (SAIL_378C01, single exon; Hilgers et al., 2018). Note that the band pattern differs in OPPP-relevant *gpt2* and *xpt* transporter mutants compared to Col wildtype and *tpt-5* (Ws wildtype corresponds to *tpt-5*, grey dashed line). Ponceau S-stained blots (protein) are shown as loading reference; RbcL, large subunit of RubisCO. Molecular masses are indicated in kDa (PageRuler Prestained Protein Ladder, Fermentas).

In addition, His-tag versions of the GPT1 and GPT2 N-termini were cloned and (following over-expression in *E. coli*) affinity-purified and used for raising polyclonal antisera in rabbits. The obtained α-GPT1 antiserum specifically recognized the N-terminus of GPT1 but not GPT2 (Supplemental Figure 14). Immunoblot analyses of different Arabidopsis tissues detected prominent high molecular weight bands in soluble fractions of flower, silique and seedling tissue - but not leaf extracts (Figure 7B), with stronger labeling in *gpt2* (Niewiadomski et al., 2005) and *xpt-2* (Hilgers et al., 2018), but not *tpt-5* mutant plants (Figure 7C). In total, four bands were found in reproductive tissues/seedlings and three bands in leaves. The latter resembled those reported for ^35^S-labeled GPT upon import into isolated plastids, namely: precursor, weak intermediate and processed *mature* forms (Kammerer et al., 1998). Intermediates are unlikely to persist *in planta*. Thus, as deduced from the stronger labeled top bands in the *gpt2* mutants compared to Col-0 wild-type, we suppose that weak bands ∼39 kDa in leaf extracts represent a minor share of active *mature* GPT1 in chloroplasts (Figure 7B, lower black arrowhead), migrating between less active *mature* (estimated 36.8 kDa) and *full-length* (estimated 42.3 kDa) versions (red arrowheads). Conversely, top bands in reproductive flower and silique tissue (black arrowheads) would represent active GPT1 in the ER/peroxisomes (Figure 7B, compare Col to *gpt2-2* and *gpt2-3*). This was also observed in seedling extracts, including other transporter mutants (Figure 7C). Interestingly, the pattern of triose-phosphate/phosphate translocator mutant *tpt-5* resembled wild-type (Ws, Col), whereas unprocessed (top) bands persisted in 4 week-old seedlings of OPPP-relevant mutants *xpt-2* and *gpt2-3*. However, additional treatments prior to SDS-PAGE/immuno-detection (-/+Lambda Protein Phosphatase, extraction -/+ phosphatase inhibitors; Supplemental Figure 14 panels F-G) or use of 200 mM DTT for tissue extraction and sample boiling (not shown), did not result in visible differences.

### GPT1 is required both at plastids and peroxisomes during fertilization

Loss of the last OPPP step in peroxisomes prevented formation of homozygous offspring due to mutual sterility of the *pgd2* gametophytes (Hölscher et al., 2016). In analogy to this, we set out to rescue plastidial versus ER/peroxisomal defects by ectopic *GPT* expression in heterozygous *gpt1* lines. First, the coding sequence of GPT2 was placed under control of the constitutive mannopine synthase (*MAS*) promoter (Guevara-Garcia et al., 1993) or the *GPT1* promoter (position −1958 to −1), and introduced into heterozygous *gpt1* plants by floral dip transformation. The CaMV-*35S* promoter-driven *GFP-GPT1*_C-mat construct (targeting the ER/peroxisomes, Figure 6C), was included for comparison (Supplemental Figure 15A). Obtained data showed that ectopic *GPT2* expression merely rescued the *gpt1* defect of incompletely filled siliques (Supplemental Figure 15B, panels a, b and f). When driven by the *GPT1* promoter, some siliques of the *ProGPT1:GPT2* transformed plants were completely filled with seeds (Supplemental Figure 15B, panel d), whereas most siliques of the same plant/line showed erratic seed maturation (panel c) or seed abortion (panel e). The frequencies of unfertilized, aborted ovules are compiled in Table 2. Compared to the untransformed heterozygous *gpt1-1* or *gpt1-2* lines (∼30%), a slight reduction was found for ::*ProMAS:GPT2* (∼27%), compared to ::*ProGPT1:GPT2* (∼21%) and Ws wild-type (∼7%), indicating some compensation by GPT2 on the female side. Attempted ER/peroxisomal rescue by ::*Pro35S:GFP-GPT1*_C-mat scored the highest values with ∼34% aborted ovules.

**Table 2.**
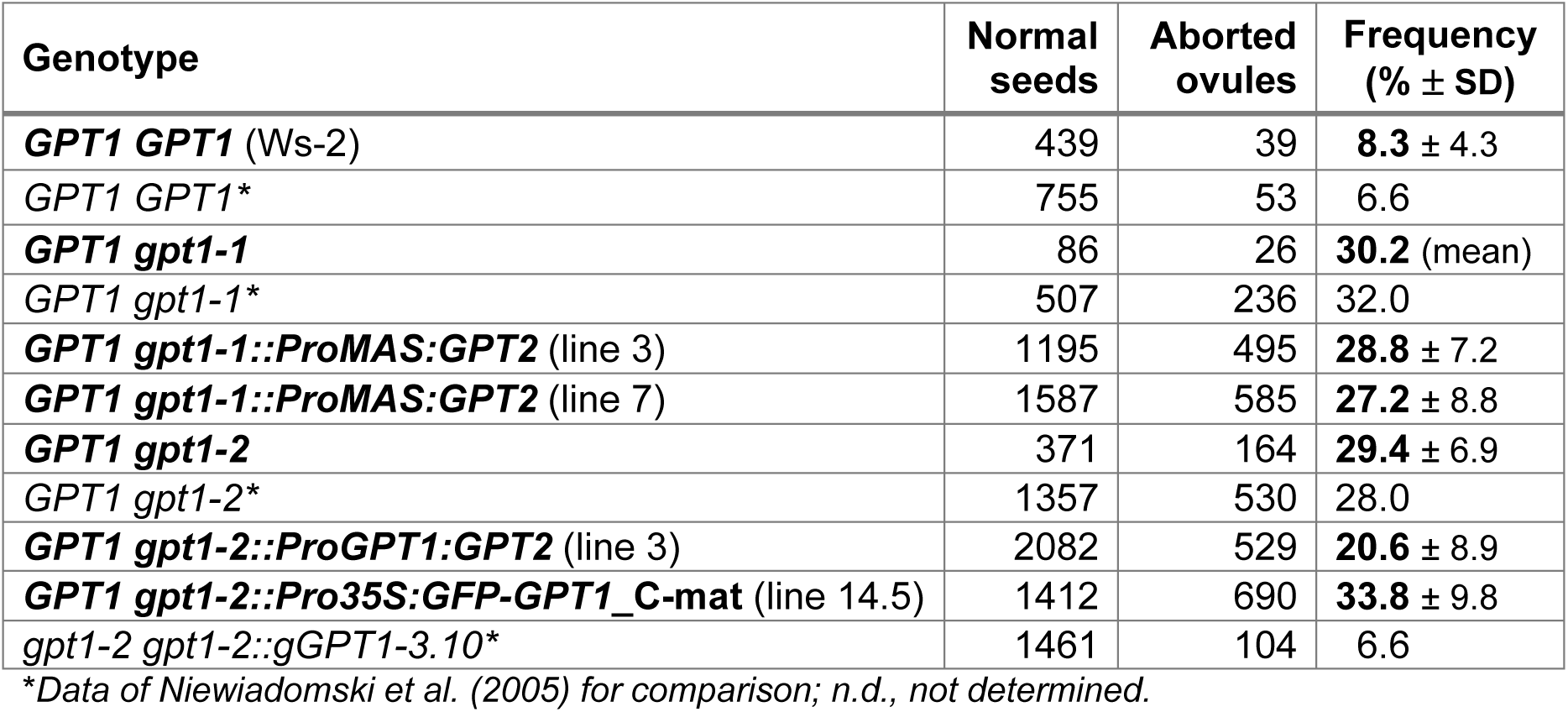
Seeds and aborted ovules without and upon ectopic GPT expression. Arabidopsis thaliana ecotype Ws-2 and heterozygous *gpt1-1* and *gpt1-2* T-DNA lines compared to plastid compensated *GPT1 gpt1-2*::*ProMAS:GPT2* or ::*ProGPT1:GPT2* lines (T2 generation), and ER/peroxisomal compensated line ::*Pro35S:GFP-GPT1*_C-mat (T3 generation). Transformed progeny was initially selected on Hygromycin B. SD, standard deviation.

Despite occasionally filled siliques, analyses of the *ProGPT1:GPT2*-compensated lines revealed no *gpt1* homozygous plants (Table 3). Therefore, *GPT1 gpt1-2*::*ProGPT1:GPT2* was reciprocally crossed with ER/peroxisomal *GPT1 gpt1-2*::*Pro35S:GFP-GPT1*_*C-mat*, forming seeds only with ::*ProGPT1:GPT2* as mother plant (Table 3). Since again no homozygous *gpt1-2* alleles were found in the F2, several T2 plants of ::*ProGPT1:GPT2* (line 3 #6 with ∼73% filled siliques; Supplemental Figure 16A) were super-transformed with *ProGPT1:GPT1*_N-long mat (ER/peroxisomal construct driven by the *GPT1* promoter; Supplemental Figure 16B), based on OFP-Pex16 co-expression (Figure 5C) and GPT1-*ro*GFP analyses (Supplemental Figure 12), but lacking the reporter.

**Table 3.** Transmission of the *gpt1* alleles with and without ectopic *GPT* expression. Segregation analysis of heterozygous *gpt1-1* and *gpt1-2* lines upon selfing or transformation with the indicated *GPT* rescue constructs: *GPT2* cDNA was driven either by the constitutive *ProMAS* (T2 generation) or the *GPT1* promoter (T2 and T3 generation). ER/peroxisomal *Pro35S:GFP-GPT1*_C-mat was analyzed in parallel (transformed plants were selected on Hygromycin). No homozygous *gpt1* plants were found. Therefore plastid-compensated *GPT1 gpt1-2*::*ProGPT1:GPT2* was reciprocally crossed with ER/peroxisomal rescue construct *GPT1 gpt1-2*::*Pro35S:GFP-GPT1*_C-mat. Only one combination set seeds, indicating that GPT2 is unable to rescue GPT1 function during pollen maturation. Still no homozygous *gpt1* plants were found. Thus, *GPT1 gpt1-2*::*ProGPT1:GPT2* was super-transformed with ER/peroxisomal rescue construct *ProGPT1:GPT1*_N-long mat (lacking the TP region) and selected on Kanamycin. Among the progeny of individuals carrying all three T-DNA alleles, *gpt1-2* transmission markedly improved, although no homozygous plants were found. Of note, this was also true for lines devoid of *ProGPT1:GPT2*. Values are given in percent with number (n) of plants analyzed.

| Genotype | *GPT1 GPT1*<br>+/+ | *GPT1 gpt1*<br>+/- | *gpt1 gpt1*<br>-/- |
| --- | --- | --- | --- |
| *GPT1 gpt1-1* | **79.3**<br>(wt = 184) | **20.7**<br>(he = 48) | **0**<br>(n = 232) |
| *GPT1 gpt1-1::ProMAS:GPT2* (lines 3 & 7, T2) | **67.8**<br>(wt = 214) | **32.2**<br>(he = 102) | **0**<br>(n = 316) |
| *GPT1 gpt1-2* | **74.8**<br>(wt = 95) | **25.2**<br>(he = 32) | **0**<br>(n = 127) |
| *GPT1 gpt1-2::ProGPT1:GPT2* (line 3, T2) | **71.0**<br>(wt = 115) | **29.0**<br>(he = 47) | **0**<br>(n = 162) |
| *GPT1 gpt1-2::Pro35S:GFP-GPT1_C-mat* (T3) | **65.8**<br>(wt = 100) | **34.2**<br>(he = 51) | **0**<br>(n = 151) |
| *GPT1 gpt1-2::ProGPT1:GPT2* (♀) x<br>*GPT1 gpt1-2::Pro35S:GFP-GPT1_C-mat* (F2)* | **80.0**<br>(wt = 152) | **20.0**<br>(he = 38) | **0**<br>(n = 190) |
| *GPT1 gpt1-2::ProGPT1:GPT2* (line 3, T3)<br>*::ProGPT1:GPT1_N-long mat* (T2)* | **56.1**<br>(wt = 184) | **43.9**<br>(he = 144) | **0**<br>(n = 328) |
| *GPT1 gpt1-2::ProGPT1:GPT1_N-long mat* (T2) | **56.5**<br>(wt = 48) | **43.5**<br>(he = 37) | **0**<br>(n = 85) |

\*progeny of plants containing all three T-DNAs; wt, wildtype; he, heterozygous; n, number analyzed; n.d., not determined

Surprisingly, siliques of heterozygous *gpt1* plants carrying *ProGPT1:GPT1*_N-long mat (T1) were almost completely filled with seeds, irrespective of whether plastidial *ProGPT1:GPT2* was present or not (Supplemental Figure 16C, compare top to bottom panels). This indicated a major contribution by GPT1 in the ER/peroxisomes, as also corroborated by the *gpt1* transmission rates (Table 3).

In summary, compared to the untransformed *GPT1 gpt1* lines (21-25%), heterozygous progeny raised only slightly upon presence of *ProGPT1*-driven *GPT2* (29-32%), with highest values scored for a *GPT1* construct lacking the transit peptide region (43%). Thus, substantial recovery by GPT1 (solely targeting the ER/peroxisomes) was obtained without further contribution by GPT2 (solely targeting plastids), expressed from the same promoter.

## DISCUSSION

### GPT1 and GPT2 differ in several aspects

Based on the concept that peroxisomes developed from the proto-endomembrane system of the Archaebacterial host in an early pre-eukaryote (Tabak et al., 2006; Cavalier-Smith, 2009; van der Zand et al., 2010), and GPT1 developed a special role related to NADPH provision by the OPPP in plastids during land plant evolution (Niewiadomski et al., 2005; Andriotis et al., 2010), as opposed to GPT2 mainly contributing to starch biosynthesis (Athanasiou et al., 2010; Kunz et al., 2010; Dyson et al., 2015), a preexisting role of GPT transporters in the secretory system is conceivable. Further support for functional specialization is reflected by the late split of GPT1 from GPT2 sequences in dicots (Figure 8), and dichotomy of orthologous sequences in the monocot species rice (*Oryza sativa*) and maize (*Zea maize*). In rice, ADP-Glc and not G6P was shown to be imported by heterotrophic plastids as the precursor of starch biosynthesis (Cakir et al., 2016), except for pollen tissue that imports G6P (Lee et al., 2016). Furthermore, the GPT1-interacting oxidoreductase Grx*_c1_* (Supplemental Table 1, listed by the MIND database also as interaction partner of GPT2, albeit with lower score) is dicot-specific, while Grx*_c2_* is present in all seed plants (Riondet et al., 2012; Li, 2014). In Arabidopsis, *GPT2* is predominately expressed in heterotrophic tissues, whereas *GPT1* is found ubiquitously (Niewiadomski et al., 2005), also in leaves (Supplemental Figure 17). Thus, basal G6P exchange, needed to stabilize the Calvin cycle in chloroplasts (Sharkey and Weise, 2016), should involve GPT1 rather than GPT2, which may be additionally induced under stress, e.g. by light (Athanasiou et al., 2010; Preiser et al., 2019).

**Figure 8.**
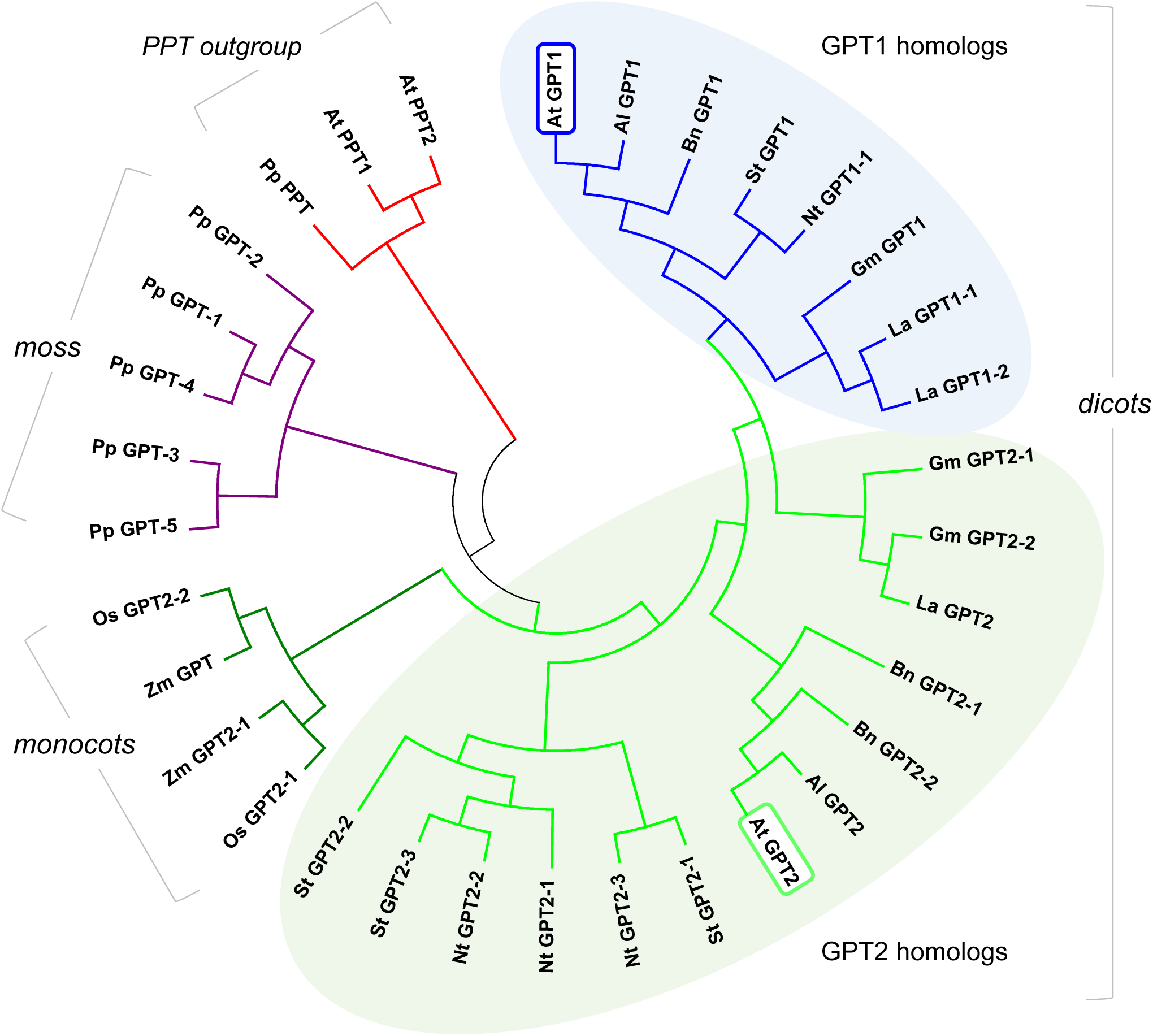
Phylogenetic analysis of GPT sequences from different plant clades. Selected GPT isoforms of the *Brassicaceae, Fabaceae, Solanaceae* and *Poaceae* in comparison to the moss *Physcomitrella patens*. The phosphoenolpyruvate/phosphate translocator (PPT) accessions serve as outgroup (red). Glucose-6-phosphate/phosphate translocators (GPT) of *Physcomitrella patens* (Pp, violet) form the base of the phylogenetic tree. GPT2 accessions (green) of monocotyledonous plants split off early (monocots, dark green), whereas the GPT1 accessions (blue) split much later from the GPT2 accessions (light green) in the dicotyledonous branch (dicots). For sequence identifications see Table S3. Abbreviations: *Al: Arabidopsis lyrata subsp. lyrata; At: Arabidopsis thaliana; Bn: Brassica napus; Gm: Glycine max; La: Lupinus angustifolius; Nt: Nicotiana tabacum; Os: Oryza sativa; St: Solanum tuberosum; Zm: Zea mays*. Evolutionary history was inferred by using the Maximum Likelihood method based on the JTT matrix-based model (Jones et al., 1992). The tree with highest log likelihood (−5414.98) is shown. Initial tree(s) for the heuristic search were obtained automatically by applying Neighbor-Join and BioNJ algorithms to a matrix of pairwise distances estimated using a JTT model, and then selecting the topology with superior log likelihood value. The tree is drawn to scale, with branch lengths measured in the number of substitutions per site. The analysis involved 34 amino acid sequences (Supplemental Table 3). All positions containing gaps and missing data were eliminated. There were a total of 252 positions in the final dataset. Evolutionary analyses were conducted in MEGA7 (Kumar et al., 2016).

### The GPT1 N-terminus mediates dual targeting

Our analyses showed that the C-terminal PTS1 motif of GPT1 is inactive, although reporter-GPT1 fusions interfered with import of the SKL-based peroxisome marker. As expected for PMPs (Rottensteiner et al., 2004), alternative GPT1 targeting was driven by other sequence motifs. The mPTS1 of class-I PMPs (directly imported into peroxisomes) comprises several positively charged amino acids on the matrix side adjacent to a transmembrane domain (Mullen and Trelease, 2006), besides a cytosolic Pex19-binding site (Rottensteiner et al., 2004; Platta and Erdmann, 2007), whereas for class-II PMPs it is only known that they exhibit an ER sorting signal (Mullen and Trelease, 2006; Eubel et al., 2008). Although the exact motif mediating ER import of GPT1 was not determined, domain swapping with GPT2 showed that the sequence must lie within the first 155 amino acids (N-terminus plus first two MDs). Since the GPT1_*N-long mat* version (without TP) was inserted into the ER, the region between K48 and the first MD (A92) is probably crucial, partly lacking and strongly differing from GPT2 (Supplemental Figure 1).

To exclude that GPT1 and GPT2 might be inserted into the ER prior to plastid import (Baslam et al., 2016) we tested Brefeldin A (BFA), a fungal toxin that inhibits the formation of ER-derived coated vesicles (Orcl et al., 1991; Klausner et al., 1992). Although BFA compartments of merged ER and Golgi vesicles were formed, GPT1 and GPT2 still localized to plastids. Furthermore, all *medial* swap constructs headed by GPT2 targeted plastids. Thus, in case of dually-targeted GPT1, threading into the plastidial Toc/Tic complex should prevent binding of the signal recognition particle (SRP) that directs proteins to the Sec61 import pore in the ER membrane (Figure 9A). Alternatively, an ER-targeting suppressor (ETS) region may be exposed by default, preventing SRP binding, as shown for human PMP70 (Sakaue et al., 2016).

**Figure 9.**
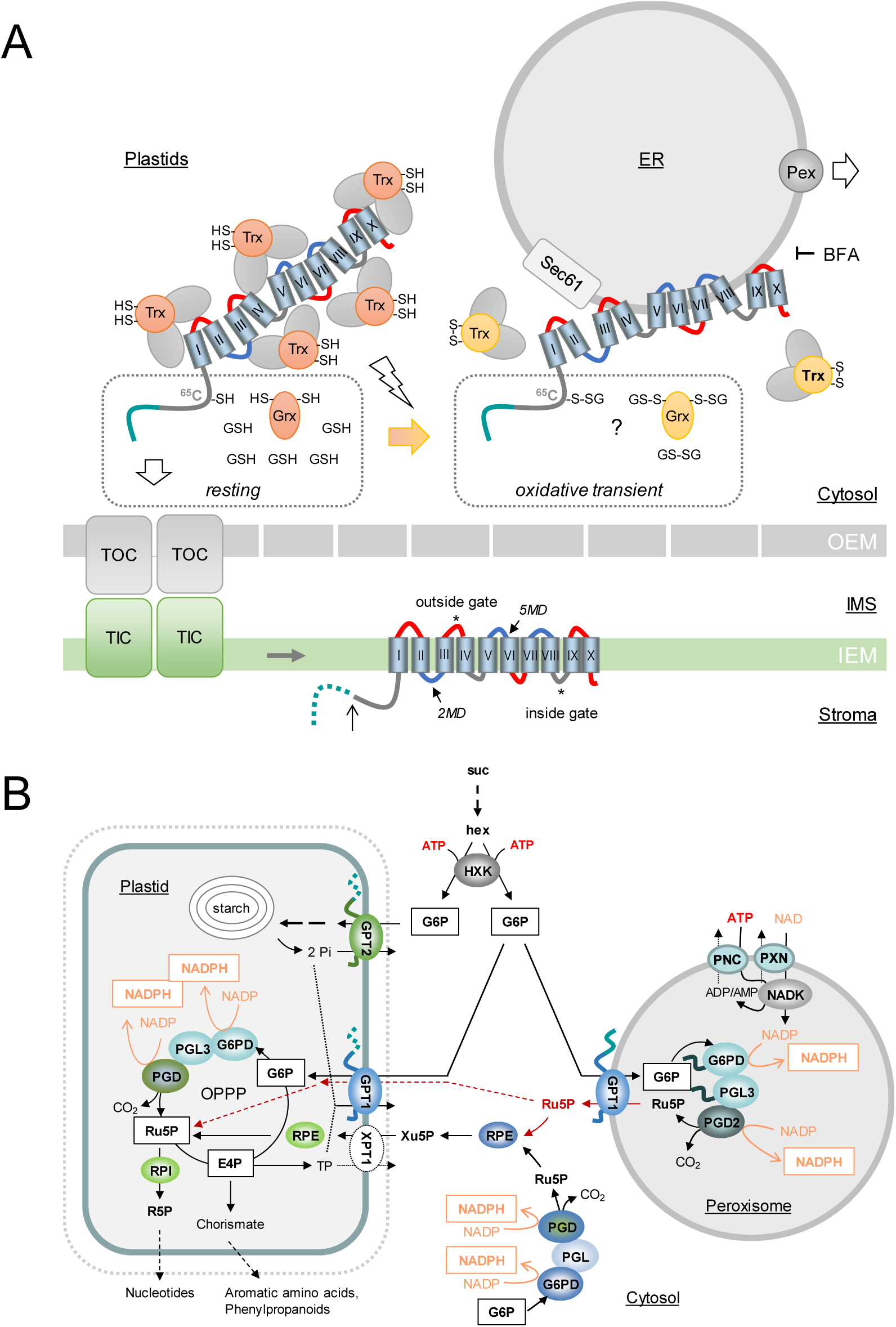
Model of dual GPT1 targeting for OPPP function in plastids and peroxisomes. **A**, GPT1 precursors in the cytosol are covered with chaperons (grey spheres) and co-chaperons Trx*_h7_* and Grx*_c1_* as putative redox sensors/transmitters (orange = reduced state, -SH; yellow = oxidized state, -S-S-). The hydrophobic membrane domains (barrels) of GPT1 are labeled with roman numerals. Hinge regions of negative net charge (blue) may facilitate ER insertion. Left, In largely reduced state of the cytosolic glutathione pool (GSH), the N-terminus of GPT1 (green) enters the TOC/TIC complex (translocon of the outer/inner chloroplast envelope), the membrane domains (MDs) integrate into the inner envelope membrane (IEM), and the transit peptide is processed (open arrow)/degraded in the stroma (dotted line). Local oxidation (flash sign) of the cytosolic glutathion pool (GSSG) likely retains GPT1 in the cytosol by a functional change in the bound redox transmitters (Grx*_c1_* and Trx*_h7_*). Whether this involves C in the GPT1 N-terminus is unclear (question mark). ER insertion involves Sec61 and sorting to peroxisomal membranes specific peroxins (Pex). Brefeldin A (BFA) blocked ER import of GPT1. **B**, Scheme of sugar metabolism in a physiological sink state. Sucrose (suc) is cleaved by cytosolic invertase yielding two hexoses (hex) that are activated by hexokinase (HXK), consuming ATP provided by glycolysis and mitochondrial respiration (not shown). By contrast to GPT2, GPT1 imports G6P into both plastids (in exchange for Pi released by GPT2-driven starch synthesis) and peroxisomes (in exchange for Ru5P that may also enter plastids via GPT1, dashed red arrows), yielding 2 moles of NADPH in the oxidative part of the OPPP. NADP inside peroxisomes is formed by NAD kinase (NADK3) that relies on ATP and NAD imported into peroxisomes via PNC (At3g05290; At5g27520) and PXN (At2g39970). The cytosolic OPPP reactions are usually linked via RPE and XPT to the complete pathway in the plastid stroma. Abbreviations: G6PD, Glucose-6-phosphate dehydrogenase; PGL, 6-Phosphogluconolactonase; PGD, 6-phosphogluconate dehydrogenase; RPE/I, ribulose-phosphate epimerase/isomerase.

How dual targeting to secretory versus endosymbiontic compartments may be regulated was discussed by Porter et al. (2015). N-terminal phosphorylation might influence competition between chloroplast import and SRP binding (as in case of protein disulfide isomerase RB60 from *Chlamydomonas reinhardtii*). GPT1 exhibits only one potentially phosphorylated serine residue in the N-terminus (S27; Supplemental Figure 1) that is conserved among all GPT sequences, albeit not listed with high score by the PhosPhAt 4.0 database (Heazlewood et al., 2008; Durek et al., 2009; Zulawski et al., 2013). Phosphomimic/preclusion of phosphorylation had no influence on dual targeting of GPT1, and neither change of the single cysteine (C65, Figure 9). On the other hand, enforced interaction of GPT1 monomers (visualized by YFP reconstitution) resulted in labeling of specific ER substructures, and the C65S change enabled detection at PerMs – as rare event (Figure 3B, panel i and 3C). However, C65 is not present in all Brassica isoforms (Supplemental Figure 1) nor in GFP-GPT1_*C-mature*, which was detected around peroxisomes upon elicitation (Figure 6C). Thus, C65 is not essential for reaching peroxisomes, but might play a role in negative regulation of GPT1 transfer from the ER to peroxisomes.

In this respect, GPT1 release to peroxisomes may require interaction with Grx*_c1_* (and Trx*_h7_*), known to engage in monothiol-dithiol mechanisms, including glutathionylation (Riondet et al., 2012; Ukuwela et al., 2018). The latter is known to be triggered by oxidative transients that accompany stress signaling and developmental change (2GSH→GSSG). Sensible cysteine residues (-S^−^ at physiological pH) may become sulfenylated (-S-OH in the presence of H_2_O_2_) or glutathionylated (-S-SG), which protects from over-oxidation (reviewed in Zaffagnini et al., 2019). Reversion (de-glutathionylation) by GSH alone is slow, but fast together with Grx and Trx (as recently shown for plastidial Amy3; Gurrieri et al., 2019). Perhaps this mechanism regulates GPT1 interaction with Pex16 and/or Pex3, given that biochemically distinct ER vesicles were shown to fuse and form new peroxisomes (Van Der Zand et al., 2012). In any case, GPT1 transport in monomeric form within the ER makes sense, since a potentially active translocator - still *en route* to its final destination - is likely not tolerated. This idea is supported by aberrant ER structure in analyses with enforced GPT1-dimer formation (Figure 3).

### Evidence for redox transmitters in GPT1 recruitment to the ER/peroxisomes

For indirect delivery of PMPs via the ER, it is still unclear how the processes of ER targeting and sorting to newly forming peroxisomes are regulated. For Pex3 it was suggested that cytosolic chaperons may guide the protein to the Sec61 translocon (Kim and Hettema, 2015), and for Pex16 that the protein may recruit Pex3 and other PMPs to the ER (Hua et al., 2015). We already published on the importance of thioredoxins as redox-dependent targeting regulators for OPPP enzymes before. Since Trx co-chaperon function (*holdase* versus *foldase*) depends on the local redox state, dual targeting of Arabidopsis G6PD1 and PGL3 is regulated by either preventing folding, allowing plastid import, or supporting folding, as pre-requisite for peroxisome import (Meyer et al., 2011; Hölscher et al., 2014). Here we show that co-expression of GPT1 with the cytosolic oxidoreductases Trx*_h7_* or Grx*_c1_* enhanced ER localization. Moreover, GPT1 interaction with both oxidoreductases was spotted at structures reminiscent of PerMs.

Thioredoxins and glutaredoxins were previously reported to promote protein folding directly, via protein-disulfide reduction or disulfide-bond formation (Berndt et al., 2008), besides enhancing co-chaperon activities in a redox state-dependent manner. Both, *foldase* function of the monomeric thioredoxin and *holdase* function in the oligomeric state, prevented folding/aggregation of client proteins, as demonstrated for Trx *h* and *m* types (Park et al., 2009; Sanz-Barrio et al., 2012). The oligomerization state of Grx*_c1_* was also shown to be influenced by the surrounding redox medium, and conversely activated under oxidizing conditions, implying a function as cytosolic redox sensor (Riondet et al., 2012; Ströher and Millar, 2012). Considering that Grx and Trx serve as electron donors for peroxiredoxins that detoxify H_2_O_2_ directly (Dietz, 2011), and regulation of *h*-type Trx via Grx*_C1_* was demonstrated previously (Meng et al., 2010; Rouhier, 2010), a complex co-regulation of the two protein classes exists in plant cells. Furthermore, Trx*_h7_* and Grx*_c1_* were found to be N-myristoylated *in planta* (Meng et al., 2010; Riondet et al., 2012; Traverso et al., 2013; Majeran et al., 2018). For Grx*_c1_*, which had been detected in the cytosol and nucleus before (Riondet et al., 2012), our results show that the protein partially resides at the ER. Grx*_c1_* promoted ER targeting of GPT1, also without N-myristoylation motif (G2A) in *grx_c1_* mutant protoplasts (not shown), indicating functional redundancy with (an)other isoform/member(s) of the Grx/Trx superfamily. Interestingly, GPT1 is listed as palmitoylation candidate by the plant membrane protein database Aramemnon (http://aramemnon.uni-koeln.de) with high score. Protein S-acylation (via cysteine residues) is still a poorly understood posttranslational process that is usually preceded by N-myristoylation, to promote membrane association, targeting, and/or partitioning into membrane subdomains (Aicart-Ramos et al., 2011; Hemsley, 2015). A potential role of Grx/Trx N-myristoylation for putative S-palmitoylation of GPT1 will have to be analyzed by a complex experimental setup, a difficult task considering partial redundancy among cytosolic Trx *h2*, *h7*, *h8*, *h9* as well as Grx *c1* and *c2* isoforms (Riondet et al., 2012; Traverso et al., 2013; Majeran et al., 2018). Clearly, GPT1 is inserted into the ER membrane in monomeric form, and may be modified at C65 (Figure 9A, question mark) for retention. Dimer formation beyond the perER would occur after de-protection, likely triggered by cytosolic redox signaling that accompanies a/biotic stress responses (Vandenabeele et al., 2004; Foyer et al., 2009) or specific developmental stages, like pollen tube elongation (Considine and Foyer, 2014) and navigation to ovules (Hölscher et al., 2016).

### GPT1 behaves like a class-II PMP

Our BiFC data suggested that GPT1 contacts at least two of the three early peroxins (Kim and Mullen, 2013). Interaction with Pex3 and Pex16 was detected at the ER and PerMs, whereas interaction with Pex19 was mostly distributed across the cytosol, reflecting its function as cytosolic cargo receptor (Hadden et al., 2006). Since simple co-expression with Pex19-reporter fusions did not show any change in GPT1 localization, dot-like structures labeled by GPT1-Pex19 BiFC analyses might be a false-positive result. This would be in line with Pex19 being mainly involved in targeting of class I, but not class II PMPs. Focal localization of GPT1 at the ER, previously described for Pex3 in yeast and for pxAPX in cottonseed/APX3 in Arabidopsis (Lisenbee et al., 2003; Narendra et al., 2006), was mainly seen upon BiFC, indicating that dimerization occurs beyond the perER. GPT1 dimers may therefore represent a forced interaction at the ER, which does not (yet) occur under physiological conditions. As a side note, Pex3 of plant cells had not been detected at the ER before (Hunt and Trelease, 2004).

Usually, GPT1 distributed evenly across the ER, unless co-expressed with Pex16 that coexists at both the ER and PerMs (Lin et al., 2004; Karnik and Trelease, 2005). Interestingly, presence of Pex16 influenced GPT1 localization at the ER, resulting in a similar but distinct pattern – also when driven by the own promoter (dark incubation in the presence of sugars activates *GPT1* mRNA expression, Supplemental Figure 18). Considering that BiFC is not dynamic, and fluorescent signals persist once the split YFP halves are reconstituted (Robida and Kerppola, 2009), GPT1 was likely dragged to PerMs upon (otherwise transient) interaction with the peroxins. In any case, this demonstrated that GPT1 can reach PerMs (although not detected there, unless triggered), wherefore the transporter may first interact with Pex16 (for ER insertion/transport to the perER; Hua et al., 2015), and then Pex3 (and possibly Pex19, during sorting to PerMs). By contrast to APX3, GPT1 is only needed at peroxisomes when the OPPP is required (Meyer et al., 2011; Hölscher et al., 2014; Lansing et al., 2019). Of note, aside from continuously imported PGD2, no other OPPP enzyme has been found by peroxisomal proteomics so far (see Hölscher et al., 2016; Lansing et al., 2019 and references cited therein).

### GPT1 transport preference differs from GPT2

After plastid import, TP sequences are cleaved off by the essential stromal processing peptidase (SPP), which is usually important for maturation, stabilization, and activation of the proteins (van Wijk, 2015). Here we show that also unprocessed GPT1 is an active transporter. Addition of a small tag or large reporter did not influence transport activity. Furthermore, topology analyses of *ro*GFP fusions indicated that upon ER insertion, both N- and C-termini of GPT1 face the cytosol (Supplemental Figure 12), similar to Arabidopsis PMP22 (Murphy et al., 2003) and the human glucose transporter (Mueckler and Lodish, 1986). These findings support the theory of Shao and Hegde (2011) that during post-translational ER import of membrane proteins, type-I topology (N-terminus facing the lumen) is strongly disfavored. This leads to obligate type-II topology (N-terminus facing the cytosol), and integration of the following MDs owing to the ‘positive inside rule’ (von Heijne, 1986; Goder et al., 2004) for the cytosolic hinge regions. The latter is not entirely true for the GPT proteins (marked red in Supplemental Figure 1 and the topology models), which may facilitate posttranslational ER insertion.

The phosphate translocator family is known to form dimers that mediate strict counter-exchange of various phosphorylated metabolites with inorganic phosphate (Pi). The ability to transport other OPPP intermediates, although possible (e.g. triose-phosphates), is usually disfavored due to the prevailing metabolite concentrations or competition with the preferred substrate (Flügge, 1999; Eicks et al., 2002). Here we show that GPT1 and GPT2 can exchange G6P for Ru5P, but GPT1 has a stronger preference for Ru5P. Thus, import of the OPPP substrate and export of its product is warranted across PerMs (Figure 9B). Moreover, poor rates obtained with 6-phospho-gluconate (6PG) as counter-exchange substrate strongly suggest that sugar-derived NADPH production occurs by all three OPPP steps (Meyer et al., 2011; Hölscher et al., 2014; Lansing et al., 2019), making a short-cut via solely Arabidopsis PGD2, catalyzing the last OPPP step in peroxisomes (Fernández-Fernández and Corpas, 2016; Hölscher et al., 2016), unlikely.

In principle, the discovered transport preference should also apply to metabolite exchange at plastids. This may explain why Arabidopsis *tpt xpt* double mutants are viable (although strongly growth-compromised; Hilgers et al., 2018) and why *rpi2* mutants, lacking one of the two cytosolic ribose-phosphate isomerase (RPI) isoforms form less starch in leaves (Xiong et al., 2009). Minute amounts of active GPT1 could drain G6P from chloroplasts due to preferred exchange with Ru5P, likely more abundant in *rpi2* mutants (Supplemental Figure 19). Besides G6P exchange needed to stabilize the Calvin cycle (Sharkey and Weise, 2016), this argues for a role of ubiquitously expressed GPT1, considering that GPT2 is absent from unstressed leaves (Supplemental Figure 14F). On the other hand, lower transport capacity of GPT1 compared to GPT2 is not surprising, since our data confirm a specialization of the two transporters. For GPT1’s function, flux rates are not necessarily a limiting parameter, but substrate specificity obviously is. This is in line with our complementation analyses, demonstrating that GPT2 cannot compensate for the absence of GPT1.

### Dual targeting of GPT1 is essential during fertilization

Niewiadomski et al. (2005) and Andriotis et al. (2010) found that loss of GPT1 function in plastids strongly affects pollen maturation and embryo-sac development, resulting in aberrant morphological changes. Interestingly, in plants with reduced GPT1 levels, embryo development is normal up to the globular stage, but then embryos fail to differentiate further and accumulate starch (Andriotis et al., 2010; Andriotis and Smith, 2019). According to the Arabidopsis eFP Browser (Winter et al., 2007), in this stage mRNA expression of *GPT2* is up to 3.5-fold higher than of *GPT1* (Supplemental Figure 17), which can explain the observed starch accumulation upon GPT1 loss.

In accordance with these premises, we suspected that ectopic *GPT2* expression may rescue some plastidial functions, but not all phenotypes of the mutant *gpt1* alleles, because swap constructs headed by GPT2 were never detected at the ER. For heterozygous *gpt1-2* transformed with *GPT2* (driven by the *GPT1* promoter), filled siliques with green, non-aborted embryos, and fertilized, but later aborted brownish embryos were observed. Plants homozygous for the *gpt1-2* T-DNA were absent from the progeny of this line and also from ER/peroxisomal compensated *Pro35S:GFP-GPT1_*C-mat.

Upon reciprocal crossing of these two lines, only one direction worked (Table 3), indicating that besides partial rescue of the female *gpt1* defects (showing as filled siliques), plastid-confined GPT2 was unable to fully rescue GPT1’s functions during pollen maturation/tube growth. Pollen grains appeared normal, but no homozygous *gpt1-2* plants were found among the progeny of combined complementation constructs. This suggested that the remaining defects result mainly from absence of GPT1 from plastids, due to a unique function GPT2 cannot fulfill. Furthermore, GPT1 transfer from the ER to peroxisomes might be impeded by artificial construct composition. Of note, *Pro35S:GFP-GPT1_*C-mat (transport-competent ER/PerM control) did not rescue ovule abortion (Table 2), but led to a substantial increase in heterozygous offspring compared to the parental line (Table 3). This may be even an underestimation, since the CaMV-35S promoter is not well expressed in pollen, and generally fluctuates in floral tissues (Wilkinson et al., 1997). By contrast, the *ProGPT1*-driven *GPT1_*N-long mat construct (without TP) rescued seed set and raised *gpt1* transmission up to 43%, independent of additional GPT2 in plastids. Thus, together with the pollination defect (mentioned above) and complementation by a genomic *GPT1* construct (Niewiadomski et al., 2005), our results indicate that for full rescue GPT1 is additionally needed in plastids, where the OPPP is mainly required for Ru5P provision to nucleotide biosynthesis (Figure 9B), as recently shown by Andriotis and Smith (2019).

The findings nicely support our previous analyses that loss of Ru5P formation in peroxisomes (by missing PGD2 activity; Hölscher et al., 2016) prevents homozygous offspring due to mutual sterility of the male and female *pgd2* gametophytes. Moreover, the low transport rates for 6PG and redundancy at the PGL step in Arabidopsis (Lansing et al., 2019) suggest that no other OPPP intermediate is transported across PerMs. Transport preference for Ru5P may also explain why GPT1 is indispensable in heterotrophic plastids (Figure 9B), probably accepting Pi released by GPT2-driven starch synthesis as counter-exchange substrate. Finally, dual targeting is supported by immuno-detection of unprocessed (ER/peroxisomal) GPT1 in flower/silique and seedling tissues. In the latter, a shift in the GPT1 pattern seems to reflect gradual adaptation to the photoautotrophic state. Besides, relative mobility and band intensities in wild-type versus *gpt2* (and other transporter mutants) indicates that GPT1 transport activity may be regulated by post-translational modification at both locations, perhaps phosphorylation of the mature protein part (up to 5 sites; Supplemental Figure 1, blue frames). Potential glutathionylation (300 Da) of the single cysteine in the GPT1 N-terminus (C65; Figure 9A) cannot explain the observed size shifts, rather palmitoylation (Greaves et al., 2008). Of note, S-palmitoylation is usually preceded by N-myristoylation (Wang et al., 1999), and both Grx*_c1_* and Trx*_h7_* were found to be N-myristoylated *in planta* (Majeran et al., 2018). For sure Grx isoforms are important during fertilization, since *grx_c1_ grx_c2_* double mutants exhibited a lethal phenotype early after pollination (Riondet et al., 2012). Together, this may add to the recently discovered role of palmitoylation during male and female gametogenesis in Arabidopsis (Li et al., 2019). However, a definite link of these aspects to dual targeting of GPT1 will require more detailed studies.

In summary, our data present compelling evidence for dual targeting of GPT1 to both plastids and peroxisomes. Imported G6P is converted by the oxidative OPPP part to NADPH and Ru5P, which is the preferred exchange substrate (likely at both locations), thus contributing to gametophyte and embryo development as well as pollen-tube guidance to ovules. Since the latter dominates the reproductive success, further analyses are required to determine the exact physiological context of GPT1’s presence at the ER/peroxisomes.

## MATERIALS AND METHODS

### Bioinformatics

For general information about *Arabidopsis thaliana*, the TAIR website (www.arabidopsis.org), Araport (www.araport.org/), PhosPhAt 4.0 (http://phosphat.uni-hohenheim.de/), and the National Center for Biotechnology Information (NCBI) (www.ncbi.nlm.nih.gov) were consulted. Routine analyses were performed with programs of the ExPASy proteomics server (www.expasy.ch) and Clustal Omega (www.ebi.ac.uk). For the phylogenetic tree, sequence information on different higher plant clades was retrieved from the National Center for Biotechnology Information (NCBI; www.ncbi.nlm.nih.gov), and for the moss *Physcomitrella patens* from www.cosmoss.org. Sequence alignments and phylogenetic analyses were performed in MEGA7 (Jones et al., 1992; Kumar et al., 2016) using the Maximum Likelihood method based on the JTT matrix-based model (Jones et al., 1992).

### Cloning of Fluorescent Reporter Fusions

Open reading frames of candidate genes were obtained by RT-PCR using Arabidopsis total leaf RNA as described in Hölscher et al. (2016), except for Trx*_h7_* which was amplified from genomic DNA. Appropriate oligonucleotide primers are listed in Supplemental Table 2. Reporter constructs were cloned in plant expression vectors as described before (Meyer et al., 2011; Hölscher et al., 2016) and indicated in the table below.

| **Reporter vector** | **Sites for N-terminal fusions** | **Sites for C-terminal fusions** |
| --- | --- | --- |
| pGFP2* | XbaI (SpeI), Acc65I | [not used] |
| pGFP2-ΔNcoI* | XbaI, Acc65I | [not used] |
| pOFP-ΔNcoI*<br>(pSY526) | EcoRI, NcoI | SpeI, BamHI |
| pGFP2-SDM* | XbaI, Acc65I | SpeI, BamHI |
| pG/OFP-NX* | XbaI, NcoI, Acc65I | SpeI, BamHI |
| pUC-SPYNE<br>pUC-SPYCE(M) | SpeI, Acc65I, BamHI | [not used] |
| pUC-SPYNE(R)<br>pUC-SPYCE(MR) | [not used] | SpeI, BamHI |

\*For vector details see (Meyer et al., 2011; Hölscher et al., 2016); split YFP vectors (Walter et al., 2004)

### Site-directed Mutagenesis

Single base changes, for destroying restriction sites or changing amino acids, were introduced by the Quick-Change PCR mutagenesis kit protocol (Stratagene), using the primer combinations listed in Supplemental Table 2 and Phusion^TM^ High-Fidelity DNA Polymerase (Finnzymes). All base changes were confirmed by sequencing.

### Heterologous Protein Expression in Yeast Cells

For in vitro-uptake studies, *full-length* or *mature* GPT1 and GPT2 versions were amplified with the corresponding primers from cDNA and inserted into yeast vectors pYES2 or pYES-NTa via Acc65I (KpnI)/BamHI sites (Thermo Scientific). For *full-length* GPT1, primer combinations were GPT1_Acc65I_s with GPT1+S_BamHI_as; for *mature* GPT1, GPT1_C-mat_Acc65I_s with GPT1+S_BamHI_as; and for *mature* GPT2, GPT2_C-mat_Acc65I_s with GPT2+S_BamHI_as (Supplemental Table 2). For the GFP-GPT1_*C-mat* version, PCR fragments (primers: GPT1_C-mat_SpeI_s and GPT1+S_BamHI_as) were first inserted into pGFP2-SDM via SpeI/BamHI sites, released with KpnI/BamHI, and cloned in pYES2. The resulting constructs were transformed into strain INVSc1 (MATa, his3Δ1, leu2, trp1-289, ura3-52/MATα,his3Δ1, leu2, trp1-289, ura3-52) using the lithium acetate/PEG method (Gietz and Schiestl, 2007). Yeast cells were selected on synthetic complete medium (SC-Ura; 0.67% (w/v) YNB supplemented with appropriate amino acids and bases for uracil auxotrophy and 2% (w/v) glucose as carbon source). Since protein expression is under control of the galactose-inducible promoter pGAL1, yeast cells were grown aerobically in SC-Ura supplemented with 2% (w/v) galactose for 6 h at 30°C. Harvest and enrichment of total yeast membranes without and with recombinant GPT proteins was performed according to Linka et al. (2008).

### Uptake Studies Using Proteoliposomes

Yeast membranes were reconstituted into 3% (w/v) L-α-phosphatidylcholine by a freeze-thaw-sonication procedure for in vitro-uptake studies as described in Linka et al. (2008). Proteoliposomes were either preloaded with 10 mM KPi, G6P, Ru5P, 6PG or produced without pre-loading (negative control). Counter-exchange substrate not incorporated into proteoliposomes was removed via gel filtration on Sephadex G-25M columns (GE Healthcare). Transport assays were started either by adding 0.2 mM [α-^32^P]-phosphoric acid (6,000 Ci/mmol) or 0.2 mM [^14^C]-glucose-6-phosphate (290 mCi/mmol). The uptake reaction was terminated by passing proteoliposomes over Dowex AG1-X8 anion-exchange columns. The incorporated radiolabeled compounds were analyzed by liquid scintillation counting. Time-dependent uptake data were fitted using nonlinear regression analysis based on one-phase exponential association using GraphPad Prism 5.0 software (GraphPad, www.graphpad.com). The initial uptake velocities were calculated using the equation slope = (Plateau - Y0)*k, whereas Y0 was set to 0. The values for the plateau and k were extracted from the non-linear regression analyses using a global fit from three technical replicates.

### Arabidopsis Mutants

Heterozygous *gpt1-1* and *gpt1-2* lines (Arabidopsis ecotype Wassilewskia, Ws-2) were kindly provided by Anja Schneider (LMU Munich) and analyzed via PCR amplification from genomic DNA as suggested for the two T-DNA alleles (Niewiadomski et al., 2005). All oligonucleotide primers are listed in Supplemental Table 2. For the Feldman line, primers GPT1_EcoRI_s/GPT1-R5 were used for the wild-type allele, and F-RB/GPT1-R5 (Niewiadomski et al., 2005) for the *gpt1-1* T-DNA allele. For the Arabidopsis Knockout Facility (AKF) line, primers GPT1-F3/GPT1-R3 were used for the wild-type allele, and GPT1-F3/JL-202 (Niewiadomski et al., 2005) for the *gpt1-2* T-DNA allele. To improve PCR analyses, GPT1-F3 was later replaced by primer gpt1-2_WT_s. Further mutants used were *gpt2-2* (GK-950D09), *gpt2-3* (GK-780F12), and *xpt-2* (SAIL_378C01) in the Columbia (Col) background, and *tpt-5* (FLAG_124C02) in the Ws background. Mutant plants were identified by genomic PCR using the suggested gene-specific and T-DNA-specific primer combinations (Supplemental Table 2).

### Plant Growth

Arabidopsis seeds were surface-sterilized by ethanol washes (vortexed for 5 s each in 70% EtOH, EtOH absolute, 70% EtOH), dried on sterile filter paper, and spread on sterile germination medium (0.5 Murashige & Skoog salt mixture with vitamins, pH 5.7-5.8, 0.8% agar; Duchefa, Haarlem, NL) supplemented with 1% sucrose and stratified for 2-3 days at 4°C. After propagation in growth chambers for one week (short day regime: 8 h light 21°C, 16 h dark 19°C) seedlings were transferred to sterile Magenta vessels (Sigma) and grown for 4-5 weeks until harvesting rosette leaves for protoplast isolation. Alternatively, seedlings were transferred to fertilized soil mix at the 4-leaf stage and grown in short day regime, prior to transfer to long day regime (16 h light 21°C, 8 h dark 19°C) to promote flowering. In case of tobacco (*Nicotiana tabacum* var. Xanthi), sterile apical cuttings were cultivated on MS agar supplemented with 2% sucrose. The top leaves of four week-old plants were used for protoplast isolation.

### Protoplast Transfection and Microscopy

Localization of fluorescent reporter fusions (all constructs driven by the CaMV-35S promoter, if not indicated otherwise) was determined by confocal laser scanning microscopy (CLSM) in freshly transfected mesophyll protoplasts (Meyer et al., 2011). For co-expression analyses, 25 µg of test DNA (BiFC: 20 µg of each plasmid) was pre-mixed with 5 µg of a reporter construct (20 µg in case of Pex16-OFP) prior to PEG transfection. After cultivation for 12 to 48 h at 21-25°C in the dark (without or with the drug/elicitor indicated), fluorescent signals were recorded using a Leica TCS SP5 microscope with excitation/emission wavelengths of 405 vs. 488/490-520 nm for *ro*GFP, 488/490-520 nm for GFP, 514/520-550 nm for YFP, and 561/590-620 nm for OFP (*m*RFP).

### Immunoblot analyses

Arabidopsis tissues were harvested from plants grown in soil, or seedlings growing on germination plates (1% sucrose) after different time points. Our standard protein-extraction buffer was 50 mM HEPES-NaOH pH 7.5, 2 mM sodium pyrosulfite (Na_2_S_2_O_5_), 1 mM Pefabloc SC, Protease Inhibitor Cocktail (1:100) for use with plant extracts (Sigma), and 280 mM β-mercaptoethanol (β-ME) - if not stated otherwise. Immunoblot analyses were conducted as described previously (Meyer et al., 2011; Hölscher et al. 2016; Lansing et al., 2019) using 10% separating gels with 10% glycerol. Polyclonal rabbit antisera were obtained from Eurogentec (Seraing, B), raised against the N-terminal GPT sequences (91 amino acids of GPT1 or 92 amino acids of GPT2) with His tag as antigen (His-N1, His-N2) after overexpression in *E. coli* BL21 from pET16b-based plasmids and affinity purification via Ni-NTA (Qiagen), followed by specificity tests (Supplemental Figure 14).

### GPT Constructs for Rescue Analyses

For one of the plastidial rescue lines, expression from the Mannopine synthase promoter was used (pBSK-pMAS-T35S, Supplemental Figure 20). The ORF of GPT2 was amplified from cDNA with primer combination GPT2_s_EcoRI/GPT2_as_PstI (all primers are listed in Supplemental Table 2) and inserted into pBSK-pMAS-T35S via EcoRI/PstI sites (pBSK-pMAS:GPT2). The entire expression cassette (pMAS:GPT2-T35S) was released with SalI/XbaI and inserted into binary vector pGSC1704-HygR (*ProMAS:GPT2*). For *GPT1* promoter-driven *GPT2*, the 5’ upstream sequence of *GPT1* (position −1 to −1958) was amplified from genomic DNA using Phusion^TM^ High-Fidelity DNA Polymerase (Finnzymes) and inserted blunt end into pBSK via EcoRV (orientation was confirmed by sequencing). The GPT2-T35S part was amplified with primers GPT2_NdeI_s/T35S_SalI_as from pBSK-pMAS:GPT2 and inserted downstream of the *GPT1* promoter via NdeI/SalI in pBSK. The final expression cassette *(ProGPT1:GPT2-T35S*), amplified with primers pGPT1_s/T35S_SalI_as, was digested with SalI, and inserted into pGSC1704-HygR via SnaBI/SalI sites.

For the CaMV promoter-driven *35S:GFP-GPT1*_C-mat construct, the expression cassette was released from vector pGFP2-SDM with PstI/EcoRI, the EcoRI site filled (using Klenow Fragment, Thermo Fisher) and inserted into binary vector pGSC1704-HygR via SdaI/SnaBI sites. For *GPT1_*N-long mat (also driven by the *GPT1* promoter), fragments were amplified with primers GPT1_long mat-s and G6P_peroxi_Trans_full_BamHI from existing cDNA clones upon insertion into the pGFP-NX backbone via XbaI and BamHI (removing GFP). The *GPT1* promoter was then amplified with primers P_GPT1_s and P_GPT1_as and inserted via PstI/SpeI into PstI/XbaI in the target plasmid, replacing the CaMV-*35S* promoter. The resulting cassette with *GPT1* promoter, *GPT1*_N-long mat and *NOS* terminator was then amplified via primers P_GPT1_s and NosT_as upon SalI digestion and insertion into SalI and SnaBI-opened binary vector pDE1001 (Ghent University, B).

All binary constructs were transformed into Agrobacterium strain GV2260 (Scharte et al., 2009). Floral dip transformation of heterozygous *gpt1* plants was conducted as described by Clough and Bent (1998). Seeds were selected on germination medium containing 15 µg ml^-1^ Hygromycin B (Roche) or 50 µg ml^-1^ Kanamycin (*ProGPT1-GPT1*_N-long mat) including 125 µg ml^-1^ Beta-bactyl (SmithKline Beecham), and transferred to soil at the 4-leaf stage. After three weeks, wild-type and T-DNA alleles were genotyped as described above. *ProMAS:GPT2* and *ProGPT1:GPT2* constructs were amplified from genomic DNA using primers GPT2_C-4MD_SpeI_s and T35S_SalI_as. For testing presence of *Pro35S:GFP-GPT1*_C-mat, primer combinations P35S_s and GPT1_EcoRI_as or NosT_as were used. Presence of *ProGPT1:GPT2* was detected with primers GPT2_XbaI_s and GPT2-Stop_BHI_as (discrimination between the cDNA-based complementation construct and wild-type sequence is based on size, i.e. absence or presence of introns), while *GPT1_*N-long mat was detected with primers GPT1_long mat_s and NosT_as”.

### Determination of Ovule-Abortion Frequencies

Siliques number 10 to 12 of the main inflorescence (counted from the top) were harvested and incubated in 8 M NaOH overnight. Images of bleached and unbleached siliques were recorded with transmitting light using a Leica MZ16 F stereo microscope connected to a Leica DFC420 C camera. Aborted ovules were counted and frequencies were calculated.

## Acknowledgements

The authors thank Anja Schneider (LMU Munich) for providing the heterozygous *gpt1* lines, and the Arabidopsis Stock Centers (NASC, INRA) for the other mutant lines. Andreas Meyer (INRES, University of Bonn) donated the *ro*GFP2 plasmid, and Robert Marschall (IBBP) helped with ratiometric analyses in ImageJ. Stephan Rips gave advice on CLSM imaging, Olessia Becker was involved in GPT antigen production/antibody testing, Sebastian Hassa cloned expression vector pBSK:pMAS-T35S, Lennart Doering finished the GPT1-C65S split YFP constructs, and Wiltrud Krüger helped with routine lab work. Several Bachelor students contributed by their thesis projects: Jan Wiese by ribose-phosphate epimerase (RPE) isoform analyses, Margareta Westphalen to BiFC interaction analyses, and Hinrik Plaggenborg to *ro*GFP topology analyses. This work was in part funded by the German Research Foundation (DFG) via grants SCHA 541/12 (to AvS) and LI 1781/1-3 (to NL).

## Author contributions

M-CB, HL, KF, TM, LC, and NL performed the experiments; AvS, M-CB, HL, and NL designed experiments and analyzed the data; M-CB, HL, and AvS wrote the manuscript; all authors read and approved the final version of the article.

## Conflict of interest

The authors declare no conflict of interest.

## Supplemental Data

The following supplemental materials are available online.

## Supplemental Tables

**Supplemental Table 1.** GPT1 search results of the Membrane-based Interactome Network Database (MIND).

**Supplemental Table 2.** Oligonucleotide primers used in this study.

**Supplemental Table 3.** Protein sequences used for calculating the phylogenetic tree.

## Supplemental Figures

**Supplemental Figure 1.** Alignment of GPT1 and GPT2 polypeptide sequences from six *Brassicaceae*.

**Supplemental Figure 2.** Localization of N-terminally truncated and full-length reporter-GPT fusions.

**Supplemental Figure 3.** Single channel images of Figure 1.

**Supplemental Figure 4.** Localization of C-terminally truncated GPT-reporter fusions.

**Supplemental Figure 5.** Localization of two different GPT1 and GPT2 *medial* reporter fusions.

**Supplemental Figure 6.** Brefeldin-A treatment of the *medial* GPT_*2MD:8MD* fusions.

**Supplemental Figure 7.** Single channel images of Figure 2.

**Supplemental Figure 8.** Single channel images of Figure 3.

**Supplemental Figure 9.** Trxh7 and Grx*_c1_* partially localize at the ER (together with GPT1).

**Supplemental Figure 10.** OFP fusions of Pex3-1, Pex16, Pex19-1 and co-expression with GFP-GPT1.

**Supplemental Figure 11.** Single channel images of Figure 5C (*Pro35S* vs. *ProGPT1* promoter).

**Supplemental Figure 12.** Ratiometric topology analysis of GPT1 at the ER using *ro*GFP.

**Supplemental Figure 13.** Single channel images of Figure 7A.

**Supplemental Figure 14.** Generation and test of the polyclonal rabbit GPT1 antiserum.

**Supplemental Figure 15.** Ectopic *GPT2* expression for plastidial rescue in heterozygous *gpt1* lines.

**Supplemental Figure 16.** ER/peroxisomal rescue of GPT1 function in heterozygous *gpt1-2*.

**Supplemental Figure 17.** Relative mRNA-expression levels of Arabidopsis *GPT1* and *GPT2*.

**Supplemental Figure 18.** G*P*T1 mRNA expression is induced by sucrose in low light/darkness.

**Supplemental Figure 19.** Possible consequences of G6P-Ru5P exchange by GPT1 at chloroplasts.

**Supplemental Figure 20.** Vector map of pBSK-pMAS-T35S.

